# The Ventral Attention Network Mediates Attentional Reorienting to Cross-Modal Expectancy Violations: Evidence from EEG and fMRI

**DOI:** 10.1101/2025.04.11.648474

**Authors:** Soukhin Das, Sreenivasan Meyyappan, Evelijne M. Bekker, Sharon Corina, Mingzhou Ding, George R. Mangun

## Abstract

Our daily interactions with the world are shaped by sensory expectations informed by context and prior experiences, which in turn influence how we allocate our attention. Prominent predictive coding models suggest that sensory expectancy and attention interact but disagree on the precise mechanisms. One possibility is that the Ventral Attention Network (VAN), may play a role by facilitating attentional reorienting when expectancy is violated. To test this, we employed an auditory-visual trial-by-trial cueing paradigm in three experiments integrating EEG and fMRI to investigate the VAN’s role in violations of cross-modal expectancy. Behavioral results showed faster responses to expected targets, confirming the efficacy of cue-induced expectations in orienting attention to the expected target modality. EEG analyses revealed differences in early (∼100 ms latency) event-related potentials (ERPs) to both auditory and visual stimuli when expectations were violated. Unexpected stimuli elicited significantly larger early-latency negative ERPs, across both modalities. Source localization of these ERPs and subsequent fMRI evidence revealed activation in the right VAN. Functional connectivity analyses further showed greater coupling between VAN regions and sensory cortices, with modality-specific pathways involving superior temporal gyrus (STG) for auditory and fusiform gyrus (FG) for visual targets. These findings demonstrate that expectancy violations recruit the VAN to reorient attention and resolve sensory conflict. By coordinating top-down control and bottom-up sensory input, the VAN supports adaptive responses to unexpected stimuli. This work advances our understanding of predictive processing in multisensory perception and highlights the VAN’s central role in flexible cognitive control.

## Introduction

Imagine that you are looking around for your cat in the house and are expecting to spot it. Instead, you hear it meow from under the couch or from another room before you see it. Such expectations and predictions are ubiquitous in our daily life and perception of the world. The human brain continuously forms expectations about incoming sensory information based on prior experiences and contextual cues. Expectations about sensory modalities, auditory and visual, have mostly been studied in isolation (Rahnev et al., 2011; Rungratsameetaweemana et al., 2018; Summerfield & Egner, 2016). Both auditory and visual expectations are thought to function within a hierarchical framework, where higher-order cortical areas generate predictions that are compared to sensory input received by lower-order areas (den Ouden et al., 2012; Kok et al., 2014; Rao & Ballard, 1999). Some theories—generally referred to as predictive coding theories—suggest that any mismatch between predicted and actual sensory information, “prediction errors”, trigger a neural signal that is inversely proportional to the magnitude of the error (Feuerriegel et al., 2021; Richter & de Lange, 2019; Summerfield & De Lange, 2014; Todorovic & de Lange, 2012; Walsh et al., 2020). Although, spatial and feature-based expectation have been investigated extensively in the past (Auksztulewicz & Friston, 2015; Bastos et al., 2012; Kok et al., 2014; Mayer et al., 2016; Samaha et al., 2018), there are very few studies that have investigated cross-modal expectations (see: (Altieri, 2014; Arnal et al., 2011; Spence & Santangelo, 2009; Stekelenburg & Vroomen, 2015; Summerfield et al., 2006).

In the context of cross-modal expectancy, predictions about one sensory modality are violated when information unexpectedly appears in another modality. Specifically, in auditory-visual modalities, expectations about one sensory domain (such as hearing) can be violated by another (such as vision). Therefore, the brain must update its predictions and reorient attention towards the unexpected, but task-relevant, stimuli. It has been shown that the Ventral Attention Network (VAN)—which includes the right temporoparietal junction (TPJ) and right inferior frontal gyrus (IFG)—is recruited to support reorienting between competing stimuli as a function of expectancy (Corbetta et al., 2008; Maurizio Corbetta & Gordon L. Shulman, 2002; Vossel et al., 2014). The TPJ is thought to support reorientation for attentional control (Geng & Vossel, 2013; Kim, 2014), while the IFG is responsible for inhibitory control to suppress irrelevant information (Aron et al., 2003; Hampshire et al., 2010). Functional neuroimaging (fMRI) studies show that VAN activation is heightened when behaviorally relevant expectations are violated (Asplund et al., 2010; Indovina & Macaluso, 2007; Kincade et al., 2005). For instance, Joseph et al. (2015) observed increased VAN activation in response to invalid gaze cues relative to valid ones. Additionally, VAN activity has been linked to inhibitory control in tasks requiring rapid suppression of responses to unexpected stimuli. Studies on stop-signal tasks have shown that VAN is activated to inhibit prepotent responses, illustrating its role in reorienting attention when predictions fail (Jahanshahi et al., 2015; van Belle et al., 2014).

Despite these findings, the VAN’s precise role in cross-modal predictive coding remains poorly understood, particularly, how it interacts with lower-order sensory regions where prediction errors are generated during sensory input violation. Questions remain about how the VAN resolves prediction errors arising from entirely different sensory modalities, such as when task-relevant auditory inputs contradict visual predictions, and about the temporal dynamics of the VAN’s involvement in such processes. EEG studies have shown that violations of expectations elicit enhanced event-related potentials (ERPs), particularly in the early and mid-latency windows (Czigler et al., 2004; Feuerriegel et al., 2021; Hall et al., 2018; M. F. Tang et al., 2018). These ERPs are thought to reflect the brain’s rapid switching response towards relevant information that was unexpected.

This study investigates the VAN’s role in cross-modal expectancy using a non-spatial, auditory-visual trial-by-trial cuing paradigm (e.g., Posner et al., 1980) which induced cross modal predictions. By integrating EEG and fMRI, we investigated the time course of neural activity and probed the VAN and sensory areas for expected and unexpected auditory and visual stimuli.

## Materials and Methods

### Overview

Three experiments were conducted at University of California Davis, using similar paradigms that manipulated cross-modal expectation. In Experiment 1, electro-encephalographic (EEG) data was recorded from participants in one session. In Experiment 2 (EEG) and Experiment 3 (functional magnetic resonance imaging; fMRI), data were obtained from different participants in separate EEG and fMRI sessions. Except for the minor differences in the procedures noted below, the analysis and methods were the same for the three experiments.

### Participants

In all three experiments, the healthy subjects reported normal hearing, normal or corrected-to-normal vision, and no history of neurological or psychological disorders. The studies were approved by the institutional review board (IRB) of the University of California, Davis. All participants provided informed written consent and were compensated financially.

In Experiment 1, EEG and behavioral data were recorded from 14 right-handed participants (mean age = 23.29 (3.58), age range = 19-30 years, 7 males). In Experiment 2, EEG and behavioral data were collected from 38 right-handed participants, but data from six participants were excluded due to poor task performance (N = 3, accuracy < 60%), excessive muscle artifacts (N = 2) and falling asleep (N = 1) during the blocks. Therefore, 32 participants were included in the final analysis (mean age = 24.09 (2.76), age range = 20-32, 9 males). In Experiment 3, fMRI and behavioral data was obtained from 12 right-handed healthy participants, but data from two participants were excluded from further analysis due to poor performance (performance was < 50%, N = 1) and failure to follow task instructions (N = 1). Thus, 10 subjects participate in the study (mean age = 22.3 (2.05), age range = 20-25, 3 males).

### Apparatus and Stimuli

#### Experiment 1

Figure 1A provides an overview of trial types and task parameters. Each trial began with the presentation of an attention-directing cue. White cues were presented on a gray background in the middle of a computer screen (19’ Viewsonic VX922 color monitor) for 250 msec. The cues instructed subjects to either prepare for an upcoming stimulus in either the auditory (Λ) or the visual (V) modality, or to passively view the upcoming stimulus (passive cues were diamond-shaped passive cues; ◊). After the cue appeared, a subsequent target was presented after a stimulus onset asynchrony (SOA) that was jittered from 1400-1600 msec. During these cue-target trials, 75% of the targets were validly cued (e.g., visual cue predicted visual target with 0.75 probability), whereas 25% were invalidly cued (e.g., visual cue followed by an auditory target). To compensate for differences in perceptual difficulty across modalities (e.g., Bridgeman, 1988), target duration was 50ms for auditory and 250ms for visual targets.

**Figure 1.**
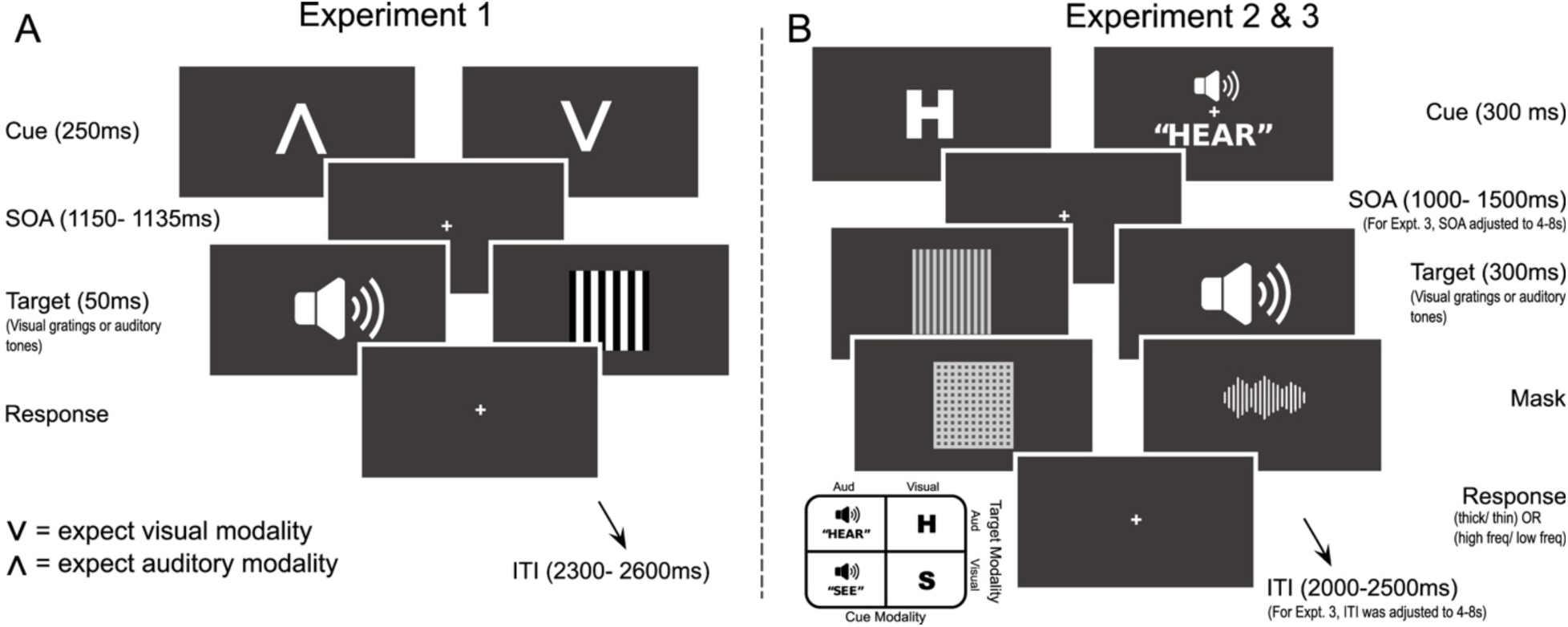
Overview of experimental design across three experiments. **(A) Experiment 1**: Participants were presented with visual cues (lasting 250 ms), either in the form of a “Λ” (expect auditory target) or “V” (expect visual target). After a variable SOA ranging between 1150-1135 ms, the target stimulus appeared, in the relevant modality (75% of the time) lasting 500 ms. The target could be either a visual (gratings) or auditory (tones) stimulus, with participants required to discriminate the frequency of the auditory tone or thickness of the visual grating. The inter-trial interval (ITI) varied randomly from 2300-2800 ms following target onset and elapsed before the start of next trial. **(B) Experiments 2 and 3**: Participants received a cue (300 ms), represented by either an “H” (“S”) or the word “HEAR” (“SEE”) to expect the modality of the upcoming target. Following an SOA of 1000-1500 ms, a target stimulus appeared for 300 ms, in the relevant modality (80% of the time). A mask was applied after the target lasting 50 ms. Participants responded by pressing buttons to indicate the type of tone or thickness of grating, The ITI in Experiment 2 was 2000-2500 ms between the onset of target and start of next trial. For Experiment 3, the SOA and ITI were both increased to 4000-8000 ms.

In the auditory modality, targets consisted of high (2000 Hz) or low (1000 Hz) pitched sine wave tones (80dB) that were presented through speakers standing on the left and right side of the computer screen. In the visual modality, targets consisted of centrally-presented black-and-white square wave gratings with a high spatial frequency of 4.8 cycles per degree (cpd) or a low spatial frequency of 0.6 cpd. Subjects were instructed to rapidly press a button with the index and middle finger of the right hand after presentation of a high tone or a high spatial frequency grating, and to press a button with the middle finger of the right hand for a low tone or a low spatial frequency grating. The mapping of stimulus attributes to response fingers was counterbalanced across subjects. Task instructions explicitly stressed that the use of information provided by the cue would increase both speed and accuracy of responding and the subjects were informed that the cues predicted the modality of the target. Each run (group of trials) included 96 cued trials (77 cue-target trials and 19 cue-only trials). In both Experiment 1 and 2 (below), the subjects were seated comfortably in a soundproof and electrically shielded room (ETS Lindgren), which was painted black to prevent any reflection of light. The room was dimly lit with DC lights to ensure a stable lighting environment.

#### Experiment 2

We utilized a non-spatial auditory-visual cue-target paradigm incorporating cross-modal probabilistic cues, as illustrated in Figure 1B. Each trial started with a 300 ms cue, which was pseudorandomly selected from one of four possible cue types: auditory verbal cues (“HEAR” and “SEE”) and visual letter cues (“H” and “S”, 0.5° x 0.5°). All auditory stimuli were pre-recorded and processed using a KEMAR recording manikin, capturing audio at the positions of the tympanic membranes with models of an average adult male pinna, and presented binaurally. Visual stimuli were presented centrally at fixation on a 24-inch ViewPixx LCD monitor positioned at a viewing distance of 850 mm. The monitor had a native resolution of 1920×1080 pixels and a refresh rate of 120 Hz (www.vpixx.com). Stimuli presentation was controlled via a Microsoft Windows 10 PC running MATLAB (MathWorks) and Psychtoolbox v3.0.8 (Pelli, 1985; Brainard, 1997). The cues probabilistically signaled the modality of an upcoming target with 80% validity. Specifically, an auditory cue (“HEAR”) or a visual cue (“H”) indicated a high likelihood that the target would be auditory, while an auditory cue (“SEE”) or a visual cue (“S”) indicated a high likelihood of a visual target. Each cue type occurred with equal probability. However, in 20% of trials, an invalid cue was presented, incorrectly predicting the upcoming target’s modality. For instance, an invalid auditory cue (“HEAR”) would be followed by a visual target instead.

During the auditory cues, the fixation cross was present on the screen. After a randomly jittered stimulus-onset-asynchrony (SOA) lasting between 1000-1500ms from cue onset, the target stimuli were presented centrally for 300ms in their respective modalities. It consisted either of a ripple tone (auditory modality) or a grating patch (visual modality). The targets were immediately followed by 50ms masks-white noise following auditory ripple tones and checkerboard stimuli of the same size after visual gratings. Participants were required to perform a two-alternative forced-choice (2AFC) task, discriminating the ripple frequency of auditory tones or the spatial frequency of visual gratings. Responses were made using two buttons, with the right index and middle fingers, and response-button mapping was counterbalanced across participants. Subjects were instructed to respond as quickly and accurately as possible within a 1000 ms response window. Each trial concluded with an inter-trial interval (ISI) jittered between 1000–1500 ms, ensuring sufficient time before the subsequent trial. Throughout the experiment, participants were required to maintain fixation at the central cross.

The auditory stimuli were created in MATLAB based on the approach by Shamma and colleagues (Shamma, 2001). Specifically, we utilized “ripples,” which are acoustic counterparts of visual gratings. We created these patterns by densely packing 100 tones that are equally spaced on the logarithmic frequency axis, ranging from 50Hz to 20kHz, with a crest-to-trough amplitude ratio of 1 and spectral peak density Ω = 0.1). By using these parameters, we generated seven tones at different ripple speeds χο = 0,2,4,8,16,20,30 Hz. As for visual targets, we generated six square wave gratings with a size of 5° x 5° and contrast of 1, having a spatial frequency of 5.1, 5.3, 5.5, 5.7, 5.9, and 6.1 cycles/degree.

To familiarize with the task and instructions, participants performed 3 blocks of a behavioral training prior to the EEG session. During this training session, participants performed all the conditions of the task until they achieved adequate performance (>90%) and proper maintenance of eye-fixation. In addition, during this phase, we used a staircase method to select two distinct tones and two distinct gratings (out of the pool) such that the participant was equally able to perform the discrimination task in both modalities. These four stimuli (two tones and two gratings) were used for the discrimination task during the main blocks. After the behavioral training session, the EEG session was divided into 8 blocks, each consisting of 80 trials (64 valid and 16 invalid trials), yielding a total of 640 trials per participant. The blocks were interspersed with a rest period of at least one minute when the meaning of the cues was emphasized, and proper eye fixation was encouraged as well.

#### Experiment 3

The task design and stimuli were identical to those of Experiment 2 (Figure 1B) but the timing was changed to permit deconvolution of overlapping BOLD responses to events that were close together in time (Das et al., 2023). We used a longer SOA of 4000-8000 ms, and longer ITI of 4000-8000 ms. The same training procedure was used outside the scanner to familiarize the participants with the task. The fMRI session was divided into 8 blocks, each consisting of 40 trials (32 valid and 8 invalid trials), totaling to 320 trials per participant.

### EEG Recording and Preprocessing

#### Experiment 1

EEG data was recorded using an elastic cap with 124 Ag/AgCl electrodes arranged in spherical coordinates (Easy cap No. M14, Falk Minow Services, Herrsching-Breitbrunn, Germany). Eye movements and blinks were recorded bipolarly from above and below the left eye (vertical electro-oculogram; VEOG), and from the outer canthi of each eye (horizontal electro-oculogram; HEOG). EEG signals were referenced to Cz during data collection. The AFz electrode (on the forehead) functioned as the ground. After task presentation, electrode positions were digitized using ELPOS (Zebris Medical GmbH, Tübingen, Germany). Data collection was continuous with a sampling rate of 250 Hz (Scan 4.3, Neuroscan Synamps 2). Online, signals were filtered with a bandpass of 0.05 Hz to 100 Hz.

#### Experiment 2

Continuous EEG data was recorded using a 64-channel Brain Products ActiCap active electrode system and digitized using a Neuroscan SynAmps2 input board and amplifier (Compumedics). Signals were recorded using Curry8 EEG acquisition program at 1000Hz without any online bandpass filtering. The 64 Ag/AgCl active electrodes were placed on the scalp using the standard international 10-10 montage, and actively referenced to a frontal-central midline electrode. To minimize any electrical interference with EEG signals, auditory stimuli were presented through earphones (ER-1, Etymotic Research, Elk Grove Village, IL, USA) via air-tubes. Additionally, electrodes at sites TP9 and TP10 were directly placed on the left and right mastoids respectively. The impedances of the electrodes were maintained below 25kΩ. Continuous data was stored in individual files corresponding to each trial block.

Signal preprocessing was carried out in MATLAB using EEGLAB toolbox (Delorme & Makeig, 2004) and Fieldtrip toolbox (Oostenveld et al., 2011). For each participant, data from every run was concatenated to form a single continuous file. Signals were visually inspected and any portion of the EEG signal containing excessive muscle artefacts, sweat potentials or large DC drifts was identified and rejected. EEG signals were re-referenced offline to the average of the left and right mastoids (TP9 & TP10). Then, EEG signals were bandpass filtered (0.1Hz-30Hz) using a Hamming window *sinc* FIR filter and then down sampled to 250Hz (for Experiment 2). Independent Component Analysis (ICA) was performed on the EEG signals to remove components that were associated with eye-blinks and eye-movements (Drisdelle et al., 2016). The ICA-processed signals for cues and targets for each trial were epoched from 200 ms before the presentation to 800 ms after their onset. Each trial was carefully screened for any eye-related or muscle-related artefacts. Furthermore, trials containing voltage exceeding +80-150 μV at any electrode location were excluded from further analysis. This process resulted in the rejection of 8.3% of trials on an average.

Subsequently, we sorted individual artifact-free EEG trials with correct responses into different experimental conditions based on type of cue (auditory “HEAR”, “SEE” and visual H and S; Λ or V), type of target based on the trial context (expected and unexpected), and modality of target (auditory or visual). Separate event-related potentials (ERP) were computed by averaging target-locked EEG epochs for each experimental condition and subject. ERP amplitudes for each experimental condition were baseline corrected using a 200 ms prestimulus window.

### fMRI Acquisition and Preprocessing

#### Experiment 3

Functional magnetic resonance images were obtained on a 3T Siemens Skyra scanner using a 20-channel head coil. Visual stimuli were presented using a MR safe VIEWPixx display (www.vpixx.com), positioned at the end of the scanner bore, and viewed via head coil mounted mirror. Auditory stimuli were presented via Sensimetric S14 insert earphones. The parameters for the EPI sequence were as follows: TR = 1800 ms; echo time = 24.4 ms; flip angle = 40°; field of view = 216 mm; 60 axial slices. The slices were oriented parallel to the plane connecting the anterior and posterior commissures. A GRAPPA acceleration factor of 2 was applied to enhance imaging speed and reduce scan time.

The preprocessing of fMRI BOLD signals was performed using the Statistical Parametric Mapping (SPM12) toolbox and custom MATLAB scripts. The preprocessing pipeline involved slice timing correction, realignment, spatial normalization, and smoothing. Slice timing correction was applied using sinc interpolation to account for variations in slice acquisition times within the EPI volume. Next, the images were realigned to the first image of each session using a 6-parameter rigid body transformation to correct for head movement during the scan. Subsequently, each participant’s images were normalized and aligned to MNI space. All images were resampled to a voxel size of 3x3x3 mm and spatially smoothed with a Gaussian kernel of 7 mm full width at half maximum (FWHM). Additionally, slow temporal drifts were removed by applying a high-pass filter with a cutoff frequency of 1/128 Hz.

### Statistical analyses

#### Behavioral Analyses

We studied participants’ behavior and how it was influenced by cross-modal expectations by calculating response accuracy and mean reaction times (RT). Trials where RT was less than 100 ms, greater than 1000 ms, or with no response were rejected (0.8% of trials). These behavioral measures were analyzed using a two-way repeated-measures analysis of variance (ANOVA) with factors expectation (expected, unexpected), and modality of target (auditory/visual). Follow-up planned two-sided paired t-tests were conducted to analyze any main effects. For all ANOVAs conducted in this study, a Greenhouse-Geisser correction was implemented in case where the assumption of sphericity was violated, and the Tukey-Kramer test was applied for post-hoc comparisons.

#### ERP Analyses (for Experiment 1 & 2, performed independently)

We analyzed the ERP components and their scalp distributions to examine how cross-modal expectations influenced target processing. First, we visually inspected the grand-averaged ERP waveforms across all participants and conditions, separately for auditory and visual targets. In both Experiments 1 and 2, this revealed a fronto-central negative ERP component peaking between 70–130 ms for auditory targets and 80–150 ms for visual targets. Past studies that have examined task-switching involving auditory and/or visual stimuli have shown similar early latency negative components, peaking between 80-120 ms for auditory stimuli and, 90-150 ms for visual stimuli (Čeponienė et al., 2003; Gajewski et al., 2018; Muller-Gass et al., 2006; Sussman et al., 2014; Vogel & Luck, 2000). Therefore, we selected these time windows to measure the ERP negativity elicited by the targets. The latency for auditory targets is faster than that of visual targets, which is consistent with the faster neural transmission and more direct cortical processing pathways of auditory information relative to visual information (Picton et al., 1974). For statistical comparisons, we used the midline electrodes and defined the scalp regions as frontal (AFz, Fz), central (Cz, CPz) and, posterior (Pz, POz) since all stimuli were presented centrally, devoid of any strong sensory hemifield bias (Li et al., 2012). To assess ERP amplitude differences, we conducted a repeated-measures ANOVA with target expectation (expected vs. unexpected) and scalp region (frontal, central, posterior) as within-subject factors. Post-hoc analyses were conducted using two-tailed paired t-tests to further explore significant effects. This set of analyses did not yield any new effects on ERP components other than the ones of interest described above.

#### Source Localization

We performed source localization on the preprocessed ERP data using standardized low-resolution electromagnetic tomography (Pascual-Marqui, 2002). First, the electrode positions were coregistered with a standard head model, and the lead field matrix was calculated. We used sLORETA to compute the cortical three-dimensional distribution of current density based on the scalp-recorded ERP data. sLORETA assumes that neighboring neurons exhibit synchronized activity and imposes smoothness constraints to solve the inverse problem. The goal of this analysis was to identify the neural sources underlying the switch-related ERPs evoked by targets under different expectation conditions (expected vs. unexpected). We focused on the same time windows where significant differences between unexpected and expected targets were found in the ERP waveforms, based on the ANOVA results.

For each participant, we computed three-dimensional sLORETA maps for both expected and unexpected targets, using averaged ERPs from the selected time windows. These maps were used to estimate the cortical sources contributing to the ERP component and to directly compare the brain activity associated with unexpected versus expected targets. At the group level, source activity between conditions (Unexpected > Expected) was compared using voxel-wise one-tailed paired t-tests. We applied a non-parametric permutation test to assess statistical significance (using 5000 permutations) and corrected for multiple comparisons using the false discovery rate (FDR) method, as implemented in sLORETA. The threshold for significance was set at p < .01. Significant source locations, where unexpected targets elicited greater activity than expected targets, were reported using Montreal Neurological Institute (MNI) coordinates. Since there were only 13 subjects in Experiment 1, due to low statistical power, the results did not survive FDR correction and reach significance. An additional goal of this analysis was to compare it with activation derived from the fMRI experiment, which used an adapted design from Experiment 2. Therefore, we report results from Experiment 2 only.

#### fMRI analyses (for Experiment 3)

We examined cue and target-evoked BOLD responses using the univariate general linear model (GLM) approach (Friston et al., 1994). Eight task-related regressors were include in the model as follows: four regressors for four types of cues (auditory “HEAR”, “SEE” & visual “H”, “S”), and four targets (expected and unexpected auditory targets, expected and unexpected visual targets) with correct responses only. We conducted a t-test by contrasting the beta values from different conditions at the voxel level to generate a t-map for each participant. We then performed group-level statistical analyses using a one-sample t-test on the t-maps from all participants. We applied a threshold of p < .05 and corrected for multiple comparisons using the false discovery rate (FDR) method in the SPM toolbox.

#### ROI definition

Bilateral IFG and TPJ were determined from the above GLM analyses to target-evoked BOLD activity. T-maps from unexpected targets were contrasted against expected targets and subjected to a p < 0.05 FDR corrected threshold. The voxels that showed statistically significant activity and were present near previously published locations were used as ROI for IFG (Weiss et al., 2018) and TPJ (Downar et al., 2000; Geng & Vossel, 2013). Furthermore, the voxels were defined by localizing them to parceled anatomical areas as defined in the AAL atlas (Rolls et al., 2020).

For functional connectivity analysis, we used a β series method to estimate the BOLD activity for each trial and voxel (Rissman et al., 2004). Each event (cues and targets) with a correct response was assigned an individual regressor. One extra regressor was included to model all the rest of the events with incorrect responses. Following this, the regressors were modeled using the GLM approach using custom MATLAB scripts within the SPM toolbox. FDR correction was applied where appropriate.

#### Functional Connectivity

We conducted a functional connectivity analysis at the single-trial level using beta (β) values (Rissman et al., 2004). Our goal was to determine whether target-related functional connectivity between regions in the Ventral Attention Network (VAN)-the inferior frontal gyrus (IFG) and the temporoparietal junction (TPJ) varied based on whether the target was expected or unexpected. At the subject level, we calculated functional connectivity by averaging β values across voxels within each region of interest (ROI) and performing a Pearson cross-correlation analysis across trials. We averaged trials based on modality (auditory and visual) and expectation (expected versus unexpected). At the group level, we derived functional connectivity by averaging each subject’s correlation coefficients, then applied Fisher’s r-to-z transformation to approximate a Gaussian distribution, which allowed us to compare connectivity across cue types.

## Results

### Behavioral Performance

In our study, participants performed a trial-by-trial auditory-visual cross-modal cuing task, where cues directed them to anticipate the modality of the upcoming target. A 2x2 repeated-measures ANOVA was used to analyze task performance and reaction times (RTs), with expectation (expected vs. unexpected) and target modality (auditory vs. visual) as factors (Figure 2, Table 1, Table 2).

**Figure 2.**
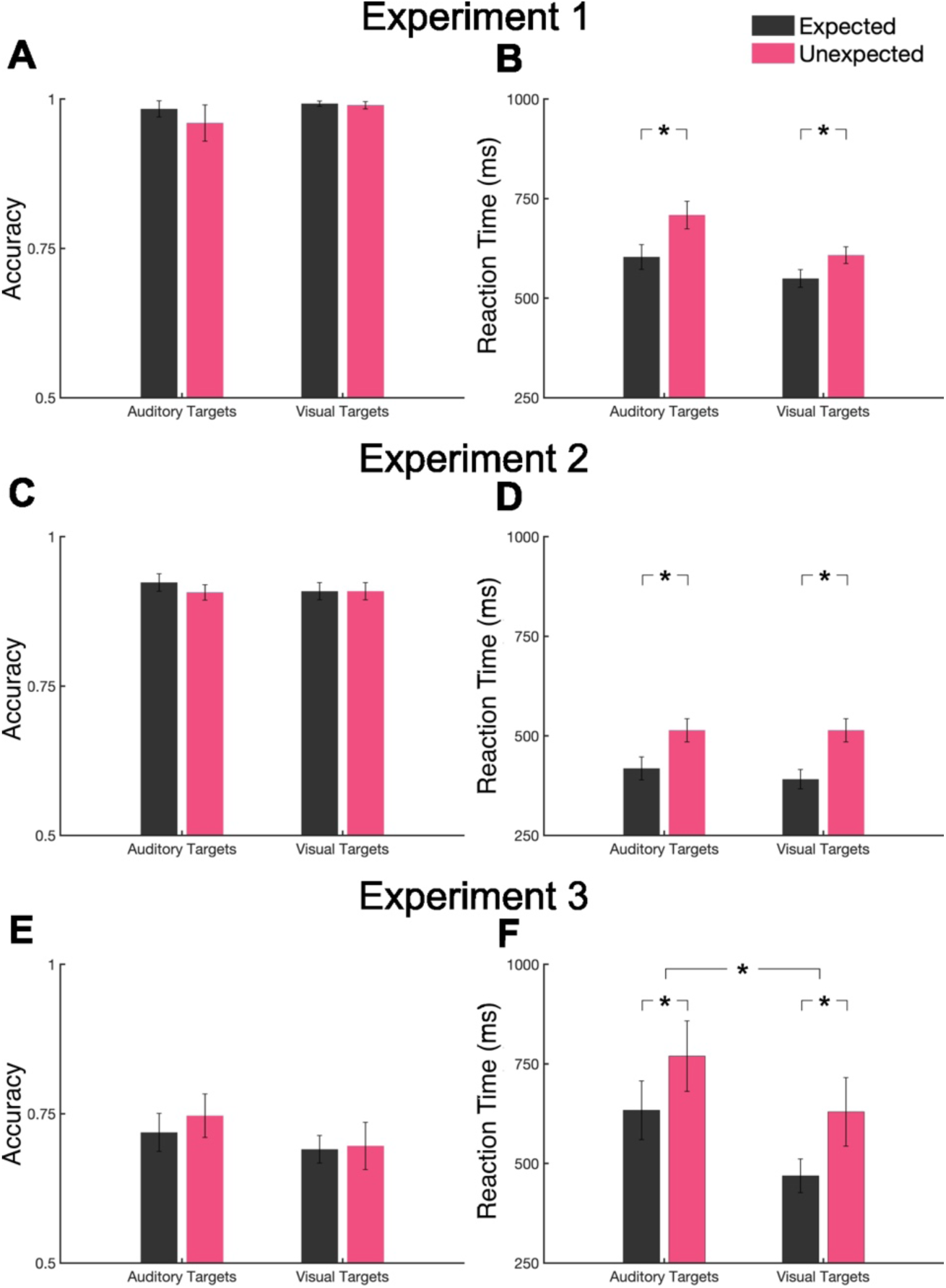
Behavioral Performance. Panels (A), (C), and (E) show participants’ accuracy in discriminating target features in Experiments 1, 2, and 3, respectively. There were no significant differences in target discrimination accuracy between auditory and visual modalities across any of the experiments. Panels (B), (D), and (F) show the reaction times (RTs) to target stimuli in Experiments 1, 2, and 3, respectively. In Panel (F), a significant main effect revealed that RTs to visual targets were faster than to auditory targets. (* denotes statistically significant main effects as indicated by ANOVA).

**Table 1.** Reaction Time (ms) for different targets across three experiments.* Values are mean (SEM). N = number of participants.

|  | N | Auditory Targets |  | Visual Targets |  |
| --- | --- | --- | --- | --- | --- |
|  |  | Expected | Unexpected | Expected | Unexpected |
| Experiment 1 | 13 | 604 (31) | 708 (34) | 549 (22) | 608 (21) |
| Experiment 2 | 30 | 418 (29) | 513 (29) | 391 (24) | 514 (29) |
| Experiment 3 | 10 | 634 (74) | 770 (88) | 469 (42) | 629 (86) |

**Table 2.** Performances for different targets across three experiments. * Values are mean (SEM).

|  | N | Auditory Targets |  | Visual Targets |  |
| --- | --- | --- | --- | --- | --- |
|  |  | Expected | Unexpected | Expected | Unexpected |
| Experiment 1 | 13 | 0.98 (.01) | 0.96 (.03) | 0.99 (.00) | 0.98 (.06) |
| Experiment 2 | 30 | 0.92 (.01) | 0.9 (.01) | 0.9 (.01) | 0.9 (.01) |
| Experiment 3 | 10 | 0.71 (.03) | 0.74 (.03) | 0.69 (.02) | 0.69 (.03) |

Across all three experiments, no significant main effect of expectation was found on task performance, indicating that the practice phase successfully balanced task difficulty across modalities. However, in each experiment, a significant main effect of expectation on RTs emerged, with participants consistently responding faster when the target stimulus was expected compared to when it was unexpected. This effect was observed in Experiment 1 (F(1,12) = 46.014, p < .001), Experiment 2 (F(1,29) = 35.688, p < .001), and Experiment 3 (F(1,9) = 8.63, p < .01), confirming that the probabilistic cues effectively shaped participants’ expectations of the target modality. In terms of target modality, differences in RTs were significant in Experiments 2 and 3, where participants responded more quickly to visual stimuli than to auditory stimuli (Expt. 2: F(1,29) = 6.094, p < .05; Expt. 3: F(1,9) = 12.135, p < .01). Interestingly, in Experiment 1, we found a significant interaction between expectation and target modality on RTs (F(1,12) = 13.47, p < .01), suggesting that the influence of expectation may have differed depending on whether the target was auditory or visual in this experiment. We did not observe any other significant interactions in Experiment 2 & 3.

Although faster responses were generally observed for visual targets, task difficulty was carefully controlled across all conditions, ensuring that participants exerted equal effort when discriminating between auditory tones and visual gratings. As a result, the focus of our EEG analyses was on within-modality differences, rather than cross-modality comparisons.

### EEG Results

#### Early ERPs to Sensory Expectations

##### Auditory Targets

In Experiment 1 (Figure 3A), a 2x2 repeated-measures ANOVA of ERP amplitude between 80-120 ms with expectation (expected vs. unexpected) and scalp regions (frontal, central, parietal) revealed a significant main effect of expectation (F(1,12) = 9.80, p < .0). Unexpected auditory targets generated more negative responses compared to expected targets. The main effect of scalp region was also significant, (F(1,12) = 50.98, p < .001), showing that the magnitude of this negative deflection was largest over fronto-central scalp regions. The interaction between expectation and scalp region was not significant (F(1,12) = 1.03, p = .33), which suggests that the effects of prediction updating were consistent across scalp regions.

**Figure 3.**
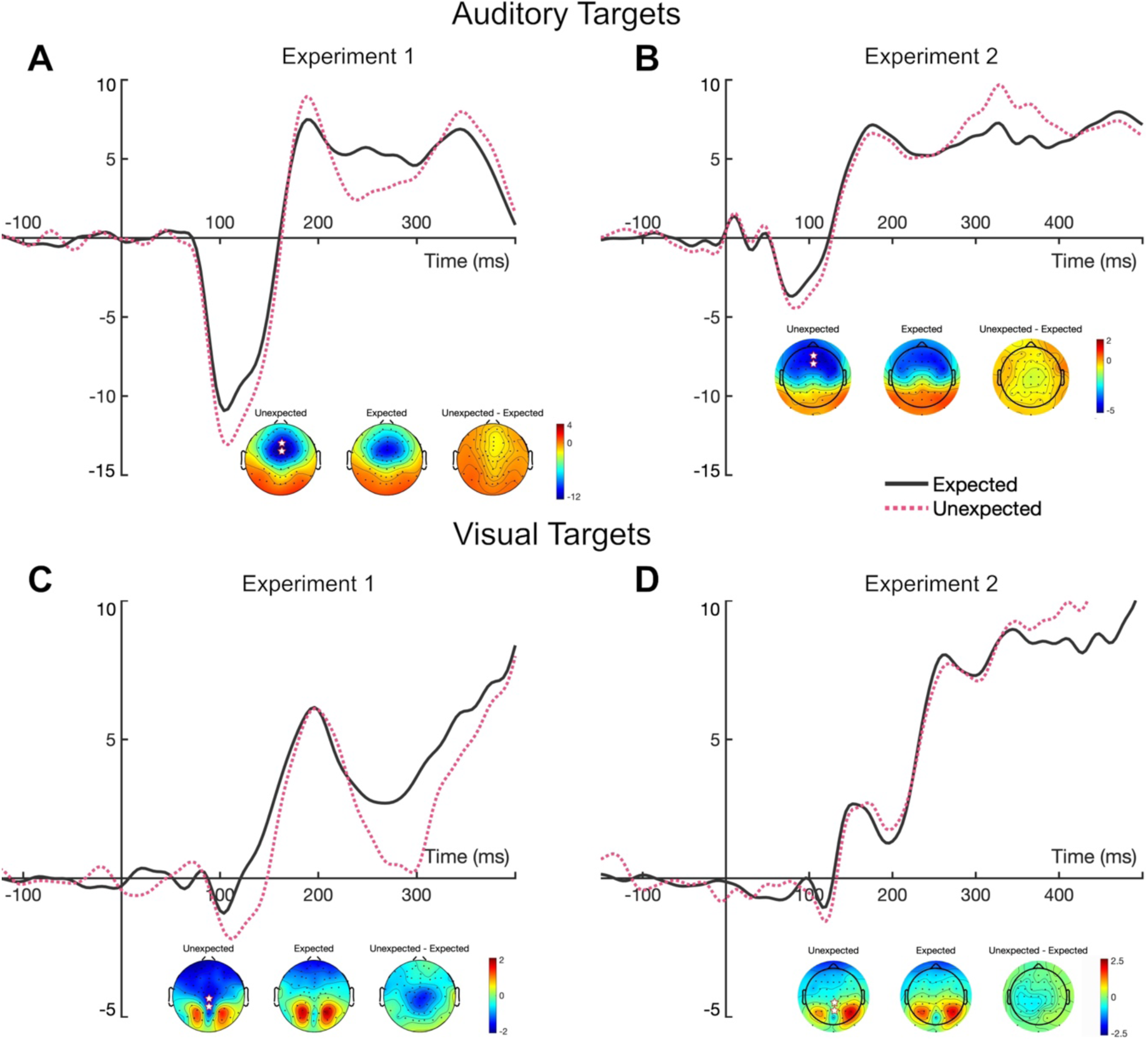
Grand-averaged ERP waveforms and topographical maps. **(A, B)** auditory target responses in Experiments 1 and 2. **(C, D)** visual target responses in Experiments 1 and 2. Expected (black) and unexpected (red) condition waveforms are overlaid. Auditory targets show greater negative deflections for unexpected stimuli, particularly over fronto-central regions. Visual targets also show greater negative deflections for unexpected stimuli, especially over central and parietal regions. Topographical maps illustrate the scalp distribution of these effects, showing stronger responses for unexpected conditions across both modalities. *Star markers in the topographical maps indicate electrodes used for generating the grand averaged ERP waveforms. Note that the target stimuli were quite different in Experiments 1 and 2, and thus the sensory-evoked waveforms have different amplitudes and slightly different latency profiles, while the negative-going deflections in the waveforms for unexpected targets is quite similar across experiments.

In Experiment 2 (Figure 3B), the ANOVA showed similar significant main effect of expectation (F(1,29) = 6.83, p < .05), with larger negative deflections for unexpected stimuli, suggesting rapid updates to the participants’ predictive models in response to violation of sensory expectations. The main effect of scalp region was also significant (F(1,29) = 22.76, p < .001), indicating differences in ERP amplitude, with the greatest deflections over fronto-central areas. Importantly, there was a significant interaction between expectation and scalp region (F(1,29) = 3.50, p < .05). Post hoc Tukey’s HSD tests showed that ERP amplitudes for unexpected stimuli were more negative in the frontal (p < .01), central (p < .05), and parietal (p < .05) regions compared to expected stimuli.

##### Visual Targets

For visual targets (Figure 3), a similar effect of switch ERP emerged during the early latency window. Unexpected visual stimuli elicited stronger negative ERP amplitudes, particularly over parietal and occipital regions divided bilaterally across the scalp, reflecting a rapid update of sensory predictions.

In Experiment 1, a 2x2 repeated-measures ANOVA on ERP amplitude with expectation and scalp region revealed a significant main effect of expectation (F(1,12) = 6.8, p < .05), indicating that unexpected visual targets elicited stronger negative ERP responses compared to expected targets (Figure 3C). However, there was no significant main effect of scalp region (F(1,12) = 0.13, p > .05) suggesting that the effects of prediction updating were not topologically specific around the scalp. Finally, the interaction between expectation and scalp region was significant (F(1,12) = 4.11, p < .05). Follow-up paired t-tests revealed stronger negativity for unexpected targets at frontal (unexpected: M = −1.70, SD = 0.10; expected: M = −0.55, SD = 0.30) and central regions (unexpected: M = −1.39, SD = 0.42; expected: M = −0.34, SD = 0.80).

Similarly, in Experiment 2, unexpected visual targets elicited significantly stronger negative deflections. The ANOVA showed a significant main effect of expectation (F(1,29) = 4.40, p < .05), reflecting the brain’s need to update visual predictions in response to unexpected targets (Figure 3D). While there was no significant main effect of scalp region (F(1,29) = 1.65, p = .21), there was a significant interaction between expectation and scalp region (F(1,29) = 3.97, p < .05). Follow-up Tukey’s HSD tests indicated that in the central region, unexpected targets (M = −0.42, SD = 0.11) produced significantly more negative ERP amplitudes than expected targets (M = 0.24, SD = 0.19, p < .05).

Despite slight differences in the experimental paradigms—such as varying cue validity (75% in Experiment 1 and 80% in Experiment 2), type of stimuli, experimental conditions, and different participant samples—the switch ERP finding was replicated across both experiments. In both cases, we observed greater negative ERP amplitudes for unexpected stimuli, suggesting that this effect is robust across different task designs and participant groups. This pattern of enhanced negativity in response to unexpected targets across both auditory and visual modalities, provides evidence for a mechanism in which the brain must rapidly updates its sensory processing when prior expectations are violated.

To summarize, across two experiment using auditory and visual cuing and target presentation, we observed a short-latency negativity elicited by unexpected modality targets. In response to auditory targets, we observed an enhanced fronto-central negativity peaking between 70 and 130 ms across both experiments. This time window corresponds to the brain’s early response to sensory input, typically associated with the auditory N1 component (Näätänen & Picton, 2007; Vogel & Luck, 2000). Similarly, we observed a centro-parietal negativity to unexpected visual targets during 90-150 ms. Traditionally, attention-driven cueing effects are characterized by larger ERPs for valid cues due to the enhanced sensory gain (Hillyard et al., 1998; Thut et al., 2006). However, in the present study, we observed what appeared to be an enhanced negativity for <u>unexpected</u> (invalidly cued) targets.

Such an effect could be considered an enhanced sensory ERP to the unexpected targets, which would align with models of perceptual expectations, where early ERP components reflect deviation from prior expectations rather than a simple attentional enhancement (Friston & Kiebel, 2009; Garrido et al., 2009; Kok et al., 2012). However, we interpret them to the result of an overlapping negativity generated in response to the requirement of switching modalities which we term as the “switch-ERP” effect. The idea is that this effect results when expectations mismatch incoming sensory input (i.e., an unexpected modality is presented), and the brain must rapidly switch to process the relevant modality, triggering a negative polarity ERP. Thus, rather than reflecting a typical cuing effect (larger ERP for valid targets or expectation induced suppression of invalid targets), the switch-ERP negativity may index additional neural resources required to shift attention to an unexpected sensory modality, aligning with prior findings on task-switching and cross-modal expectation violations (Bekker et al., 2005; Mazaheri et al., 2010). This argument is supported by the fMRI results presented below.

#### Source Localization of the Enhanced Negativity in early latency ERPs

To investigate the neural sources underlying the observed switch-ERP effect (unexpected > expected) in Experiment 2, we performed source localization on the difference wave (unexpected – expected), averaged between the pre-determined time windows using sLORETA. As a result, we observed activation in different brain areas which are recruited when processing unexpected sensory information and re-orientation to the relevant modality.

##### Auditory Targets

We found key areas in the frontal and temporal regions of the brain that were significantly activated (Figure 4A). Voxels in the Superior (BA 8/9) and Middle Frontal Gyrus (BA 10/11) exhibited bilateral activity, with more activation in the right hemisphere. The Inferior Frontal Gyrus (BA 46/47) also showed significant activation, with peak activation in the left but greater number of voxels in the right hemisphere. Moving to the posterior regions, the Precuneus (BA 7) was highly active bilaterally, with the right hemisphere showing extensive activation, suggesting a role in higher-order cognitive processes related to auditory stimuli. The Superior Parietal Lobule (BA 7) also revealed strong bilateral activation along with the Superior Temporal Gyrus (BA 22/38), again more dominant in the right hemisphere. Temporal regions, particularly the Middle Temporal Gyrus (BA 21/39) and Inferior Temporal Gyrus (BA 20), showed moderate activation bilaterally. For all activated regions and summary see Table 3.

**Figure 4.**
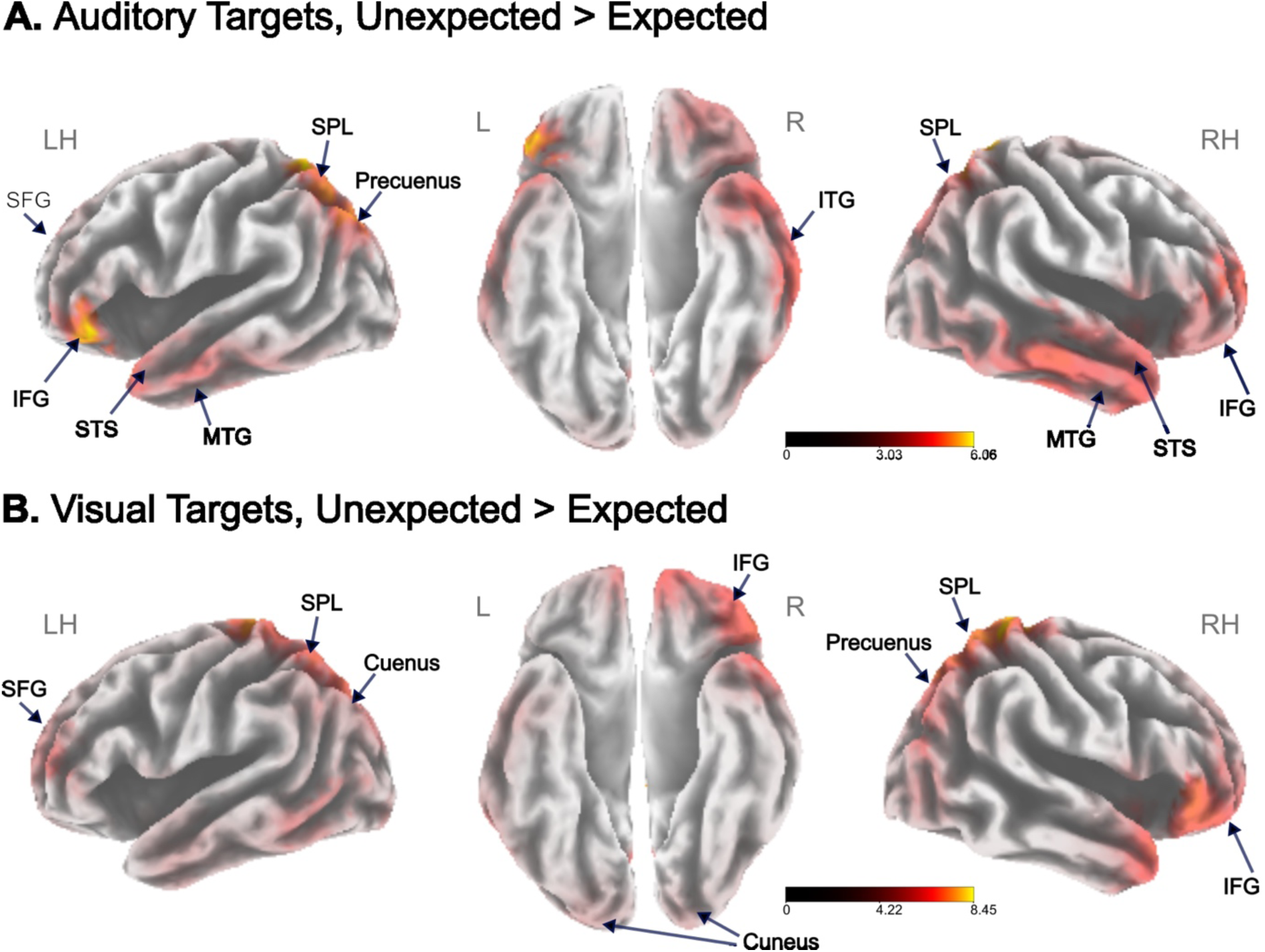
Source localization. Results of sLORETA modeling for **(A)** auditory and **(B**) visual target responses (unexpected > expected). The t-maps show cortical regions that were significantly activated during the processing of unexpected targets compared to expected targets. Source activations were processed using Statistical Non-Parametric Mapping (SnPM) and co-registered to the probabilistic MNI-152 template (Mazziotta et al., 2001). Highlighted areas in yellow and red show regions of significant activation (p < 0.01).

**Table 3.** sLORETA analysis for Auditory Targets in the 70-120 ms time window (unexpected – expected). MNI Coordinates correspond to peak activity in a brain region. BA = Brodmann area, L = Left, R = Right, MNI = Montreal Neurological Institute, Mean = mean activation of all voxels, Max. = peak activation of voxel in that region.

| Brain Region | BA | L/R | MNI Coordinates |  |  | t-value |  | No. of<br>activate<br>d voxels |
| --- | --- | --- | --- | --- | --- | --- | --- | --- |
|  |  |  | X | Y | Z | Mean | Max. |  |
| Superior Frontal Gyrus | 8/9 | L | 8 | -10 | 50 | 3.86 | 4.80 | 49 |
|  |  | R | 8 | 10 | 50 | 3.38 | 4.17 | 76 |
| Middle Frontal Gyrus | 10/11 | L | 11 | -40 | 35 | 3.74 | 5.68 | 66 |
|  |  | R | 45 | 50 | 15 | 4.02 | 5.21 | 78 |
| Inferior Frontal Gyrus | 46/47 | L | 47 | -55 | 35 | 3.99 | 6.20 | 106 |
|  |  | R | 45 | -60 | 55 | 3.69 | 4.31 | 116 |
| Precuneus | 7 | L | -25 | -75 | 50 | 4.65 | 5.88 | 113 |
|  |  | R | 10 | -80 | 45 | 4.43 | 5.62 | 119 |
| Cuneus | 17/18 | L | 19 | -5 | -90 | 4.11 | 5.22 | 68 |
|  |  | R | 5 | -90 | 35 | 3.94 | 5.41 | 78 |
| Superior Temporal Gyrus | 22/38 | L | -55 | 10 | -5 | 3.82 | 5.26 | 112 |
|  |  | R | 65 | -20 | 0 | 3.92 | 4.51 | 136 |
| Middle Temporal Gyrus | 21/39 | L | -65 | -5 | -15 | 3.72 | 4.58 | 109 |
|  |  | R | 65 | -25 | -15 | 4.17 | 4.65 | 106 |
| Inferior Temporal Gyrus | 20 | L | 65 | -30 | -20 | 3.88 | 4.73 | 54 |
|  |  | R | 65 | -30 | -20 | 3.88 | 4.73 | 49 |
| Superior Parietal Lobule | 7 | L | -20 | -60 | 65 | 5.43 | 6.33 | 65 |
|  |  | R | 20 | -60 | 65 | 4.84 | 5.98 | 63 |
| Inferior Parietal Lobule | 40 | L | 40 | -40 | -65 | 5.29 | 3.74 | 54 |
|  |  | R | 45 | -60 | 55 | 4.54 | 3.62 | 74 |
| Middle Occipital Gyrus | 18/19 | L | -20 | -100 | 5 | 3.64 | 4.80 | 118 |
|  |  | R | 20 | -100 | 5 | 3.62 | 4.66 | 55 |

##### Visual Targets

The sLORETA analysis of visual targets indicated extensive activation across different parts of the brain, overlapping with regions present in the visual processing networks. As shown in Figure 4B, brain regions in the right hemisphere were dominant. The frontal areas, including Superior (BA 8/9) Middle (BA 10/11) and Inferior Frontal Gyrus (BA 45/47) showed strong bilateral activations, indicating their role in visual reorientation and higher-order visual processing.

In posterior regions, the Precuneus (BA 7) and Cuneus (BA 17/18) demonstrated bilateral engagement, with stronger activation on the right side. Furthermore, the Superior Temporal Gyrus (BA 22/38) was involved bilaterally, but with stronger right hemisphere activation. Finally, we also observed significantly activated voxels in the Superior (BA 7) and Inferior Parietal Lobules (BA 39/40), particularly in the right hemisphere. See Table 4 for a list of activated regions and statistics.

**Table 4.** sLORETA analysis for visual targets (unexpected – expected) during the 90-130 ms time window). MNI Coordinates correspond to peak activity in a brain region. BA = Brodmann area, L = Left, R = Right, MNI = Montreal Neurological Institute, Mean = mean activation of all voxels, Max. = peak activation of voxel in that region.

| Brain Region | BA | L/R | MNI Coordinates |  |  | t-value |  | No. of<br>activate<br>d voxels |
| --- | --- | --- | --- | --- | --- | --- | --- | --- |
|  |  |  | X | Y | Z | Mean | Max. |  |
| Superior Frontal Gyrus | 8/9 | L | -35 | 55 | 20 | 4.61 | 6.34 | 181 |
|  |  | R | 15 | 65 | -15 | 4.82 | 6.71 | 200 |
| Middle Frontal Gyrus | 10/11 | L | -40 | 50 | 15 | 4.01 | 6.30 | 204 |
|  |  | R | 50 | 40 | -5 | 4.70 | 6.97 | 170 |
| Inferior Frontal Gyrus | 45/47 | L | -45 | 40 | 15 | 3.61 | 5.52 | 154 |
|  |  | R | 55 | 35 | 0 | 4.69 | 7.20 | 156 |
| Precuneus | 7 | L | -5 | -50 | 60 | 5.43 | 7.32 | 163 |
|  |  | R | 5 | -50 | 60 | 5.51 | 7.57 | 135 |
| Cuneus | 17/18 | L | -30 | -90 | 30 | 4.06 | 6.15 | 129 |
|  |  | R | 15 | -95 | 25 | 4.02 | 5.51 | 103 |
| Superior Temporal Gyrus | 22/38 | L | -60 | 0 | 5 | 4.01 | 5.08 | 168 |
|  |  | R | 50 | 20 | -20 | 4.59 | 6.89 | 177 |
| Middle Temporal Gyrus | 21/39 | L | -60 | -65 | 5 | 4.67 | 6.05 | 167 |
|  |  | R | 55 | 10 | -25 | 4.35 | 6.28 | 176 |
| Inferior Temporal Gyrus | 20 | L | -60 | -65 | -10 | 4.74 | 5.70 | 78 |
|  |  | R | 50 | -75 | -5 | 4.06 | 5.38 | 73 |
| Lingual Gyrus | 18/19 | L | -25 | -95 | -10 | 4.20 | 5.63 | 89 |
|  |  | R | 20 | -100 | -10 | 4.13 | 5.37 | 80 |
| Superior Parietal Lobule | 7 | L | -60 | -65 | -10 | 4.74 | 5.70 | 78 |
|  |  | R | 50 | -75 | -5 | 4.06 | 5.38 | 73 |
| Inferior Parietal Lobule | 39/40 | L | -35 | -55 | 60 | 4.32 | 5.96 | 117 |
|  |  | R | 40 | -70 | 45 | 4.11 | 6.00 | 116 |
| Fusiform Gyrus | 20/37 | L | -60 | -5 | -30 | 3.93 | 5.34 | 115 |
|  |  | R | 20 | -95 | -20 | 3.92 | 5.08 | 110 |

#### fMRI Results

To further investigate the enhanced ERP negativity observed when sensory expectations were violated during the early latencies, we conducted fMRI using the same stimuli and an adapted-to-fMRI experimental design (Das et al., 2023). Our primary goal was to determine whether unexpected targets evoke distinct neural activity compared to expected targets. Secondly, we aimed to identify the cortical areas responsible for processing unexpected information and relate them to the observed ERP differences. To this end, we directly compared BOLD activity between unexpected and expected targets using a general linear model (GLM) analysis.

#### fMRI Activations (Unexpected > Expected Targets)

##### Auditory Targets

The auditory fMRI analysis revealed several brain regions that showed greater activation in response to unexpected auditory targets compared to expected ones. Significant differences were observed across multiple regions, including frontal, temporal and parietal areas (see Table 5, Figure 5A-B).

**Figure 5.**
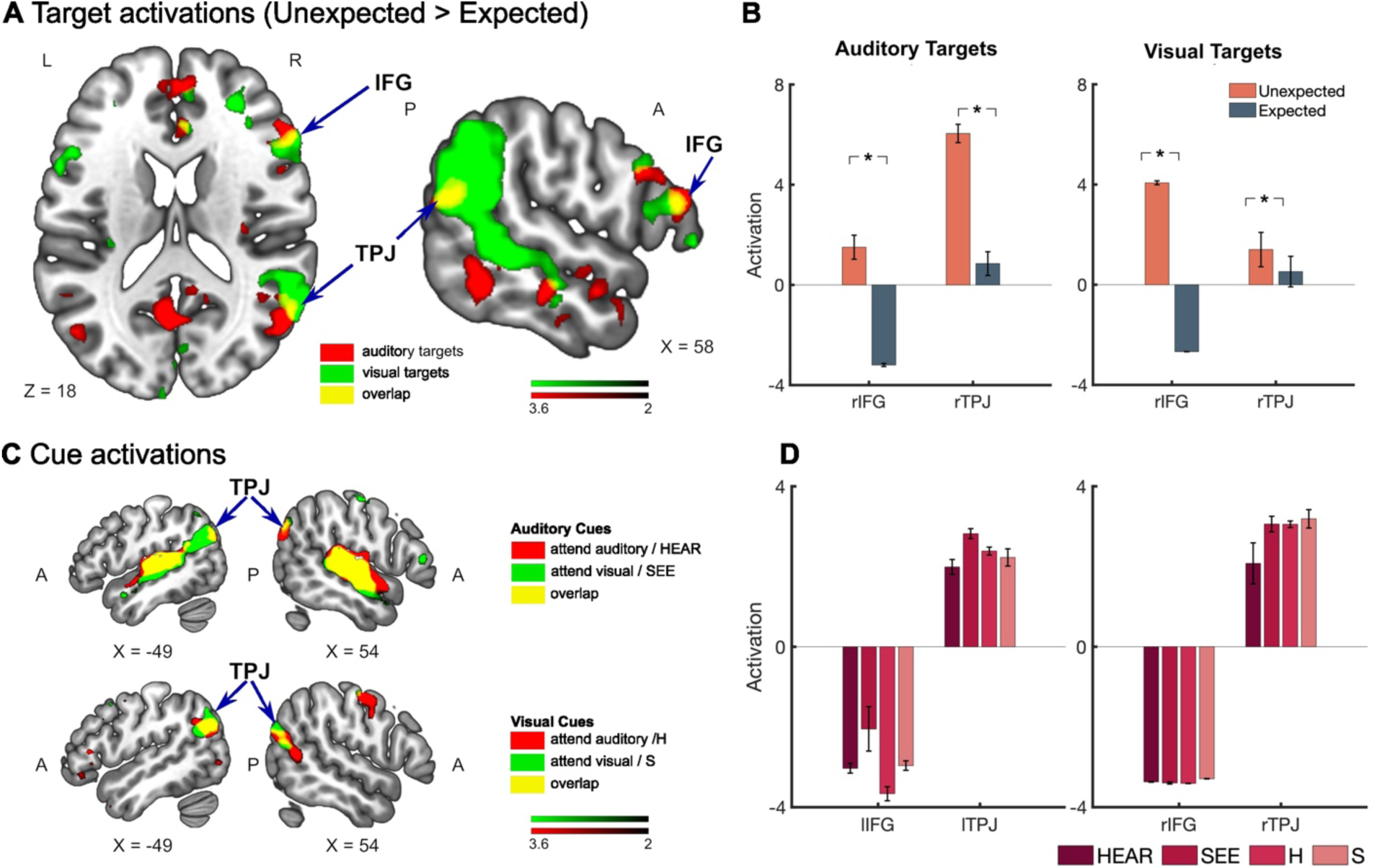
Activation of the Ventral Attention Network (VAN) in response to unexpected versus expected sensory stimuli across auditory and visual modalities. **(A)** Axial and sagittal brain slices showing significantly greater activation in the right inferior frontal gyrus (rIFG) and right temporoparietal junction (rTPJ) for unexpected stimuli compared to expected stimuli, across auditory (red) and visual (green) targets. (B) Increased activation in the rIFG and rTPJ for unexpected stimuli in both auditory and visual modalities. This suggests that the VAN, is engaged in response to cross-modal prediction violations. **(C, D)** Cue related activations in left TPJ and right TPJ for auditory (top row) and visual (bottom row) cues.

**Table 5.** MNI coordinates and corresponding t-values for brain regions that showed greater activation in response to unexpected compared to expected auditory targets. BA = Brodmann areas, L/R = Left/Right.

| Brain Region | BA | L/R | MNI Coordinates |  |  | t-value |
| --- | --- | --- | --- | --- | --- | --- |
|  |  |  | X | Y | Z |  |
| Superior Frontal Gyrus | 11 | L | -18 | 47 | 2 | 2.13 |
|  |  | R | 21 | 44 | -13 | 2.73 |
| Middle Frontal Gyrus | 45/11 | L | -33 | 47 | 17 | 2.12 |
| Inferior Frontal Gyrus | 45/47 | L | -48 | 44 | -13 | 3.30 |
|  |  | R | 39 | 56 | -10 | 1.81 |
| Superior Temporal Gyrus | 22 | L | -57 | -19 | -1 | 2.01 |
| Middle Temporal Gyrus | 21 | R | 48 | -31 | 5 | 1.94 |
| Lingual Gyrus | 17/18 | R | 6 | -79 | -4 | 3.27 |
| Supramarginal Gyrus | 6 | R | 54 | -1 | 53 | 2.40 |
| Lateral Occipital Gyrus | 19 | L | -30 | -79 | -25 | 2.28 |
|  |  | R | 48 | -76 | -16 | 3.69 |

In the frontal areas, bilateral superior, middle, and inferior frontal gyri, exhibited increased activation, suggesting the involvement of executive processes when the auditory target was not expected. Of particular interest was the increased activation of the right inferior frontal gyrus (IFG), known to play a key role in inhibitory control and the reallocation of attention to novel or unexpected events (Aron et al., 2003). Increased activity in right IFG is typically associated with suppression of expected responses so that the brain can reorient towards unexpected auditory stimuli (Hampshire et al., 2010). Additionally, in the temporal regions, superior temporal gyrus on the left hemisphere, traditionally associated with primary auditory processing, displayed increased activation, indicating its role in processing the unexpected auditory information (Yi et al., 2019). Additionally, the right middle temporal gyrus (MTG) showed greater activation, reflecting its involvement in processing complex auditory stimuli, such as semantic content or novel auditory features (Ren et al., 2020; Weiss et al., 2018). The right temporoparietal junction (TPJ), including regions surrounding the MTG and supramarginal gyrus, also demonstrated significant activation. The right TPJ is critical for bottom-up attention processes, particularly for detecting and responding to unexpected stimuli (Maurizio Corbetta & Gordon L. Shulman, 2002; Solís-Vivanco et al., 2021). Other regions, such as the lingual gyrus and lateral occipital gyrus, also showed increased activation in response to unexpected auditory targets, potentially indicating cross-modal processing triggered by unexpected events (Downar et al., 2000; Sidlauskaite et al., 2014).

##### Visual Targets

The fMRI analysis for visual targets similarly revealed stronger activation in several brain regions when participants processed unexpected visual stimuli compared to expected ones (for all regions, see Table 6). Similar to auditory targets, right inferior frontal gyrus (IFG) and right temporo-parietal junction (TPJ) were more activated for unexpected visual targets compared to expected ones (Figure 5A-B).

**Table 6.** MNI coordinates and corresponding t-values for brain regions that showed greater activation in response to unexpected compared to expected visual targets. BA = Brodmann areas, L/R = Left/Right.

| Brain Region | BA | L/R | MNI Coordinates |  |  | t-value |
| --- | --- | --- | --- | --- | --- | --- |
|  |  |  | X | Y | Z |  |
| Superior Frontal Gyrus | 11 | L | -27 | 44 | -7 | 3.47 |
| Middle Frontal Gyrus | 11 | R | 0 | 38 | -28 | 2.87 |
| Inferior Frontal Gyrus | 47 | R | 33 | 56 | -10 | 1.94 |
| Middle Temporal Gyrus | 21/36 | L | -33 | 5 | -40 | 3.56 |
|  |  | R | 27 | 14 | -43 | 2.37 |
| Inferior Temporal Gyrus | 20 | L | -54 | -7 | -34 | 4.01 |
|  |  | R | 48 | -7 | -34 | 5.64 |
| Lingual Gyrus | 18 | R | -15 | -64 | -4 | 5.30 |
| Intraparietal Sulcus | 7 | R | 18 | -61 | 53 | 2.05 |
| Intraparietal Sulcus | 40 | R | 36 | -46 | 47 | 1.94 |
| Supramarginal Gyrus | 2 | L | -63 | -25 | 35 | 3.43 |
| Fusiform Gyrus | 20/37 | L | -39 | -49 | -13 | 3.66 |
|  |  | R | 33 | -58 | -16 | 2.30 |

Additionally, in the frontal cortex, we observed increased activation in the left superior frontal gyrus and the right middle frontal gyrus. Furthermore, in the temporal areas, bilateral middle temporal gyrus (MTG) and the right inferior temporal gyrus (ITG) showed greater activation to unexpected visual targets. Importantly, significant activation was also found in the right TPJ, including regions such as the MTG, inferior parietal lobule, and lateral occipital gyrus.

Finally, we also examined cue-related activity within the same VAN voxels that were activated during target processing (Figure 5C-D). Interestingly, in contrast to the right-lateralized activations observed for targets, both left and right TPJ were activated in response to cues. However, we observed deactivation in bilateral IFG during this cue period. This is expected during the cue period, as the brain is actively forming predictions and preparing for the anticipated modality, which reduces the demand for inhibitory control typically associated with IFG activation (Braga et al., 2013; Shulman et al., 2003; Vossel et al., 2014).

#### Overlap between Source Localization and fMRI Activations

The fMRI results from Experiment 3 revealed key activations within fronto-parietal regions, indicating the engagement of VAN in response to unexpected stimuli, across both auditory and visual modalities. While these findings confirm VAN’s involvement in processing cross-modal violations, the poor temporal resolution of fMRI (Das et al., 2023) limits our ability to delineate the precise timing and dynamics of VAN. To solve this, we incorporated the source localization areas from Experiment 2, overlaying the thresholded (p < .05) sLORETA maps onto the fMRI-derived difference maps from Experiment 3. By calculating sLORETA maps of averaged difference of ERPs in successive 40 ms time windows around the peak of the switch ERP effect, we aimed to precisely identify the temporal evolution of neural responses within the VAN.

For auditory targets (Figure 6A–C), the overlap between ERP source localization and fMRI activations began to emerge during the 80–120 ms window, coinciding with the observed auditory switch ERP effect. During this window, overlap occurred over both right IFG and right TPJ, converging evidence in favor for their critical role in response inhibition that is essential for the modality switch during the unexpected stimuli. Following that, during the 120–160 ms window, the overlap reduced, remaining mainly in the right IFG. In case of visual targets (Figure 6D–F), we started the 40 ms moving window from 48 ms, so that we could capture the preselected time window (90-150 ms). Here, the overlap between ERP and fMRI activations appeared slightly earlier, with the right IFG activation beginning during the 48–88 ms window, even before the onset of the visual switch ERP effect. During the peak visual switch ERP window (88–128 ms), both right IFG and TPJ show prominent activation, paralleling the pattern seen in auditory processing. After this peak window, the overlap was confined to the right IFG only. Interestingly, in both auditory and visual modalities, the right TPJ overlap was prominent only during the peak switch ERP time window. This progression suggests that while both TPJ and IFG are involved early on, TPJ’s engagement is more transient and is related to the enhanced negativity of the switch ERP, while IFG continues to support the ongoing inhibitory control required to manage the task switch.

**Figure 6.**
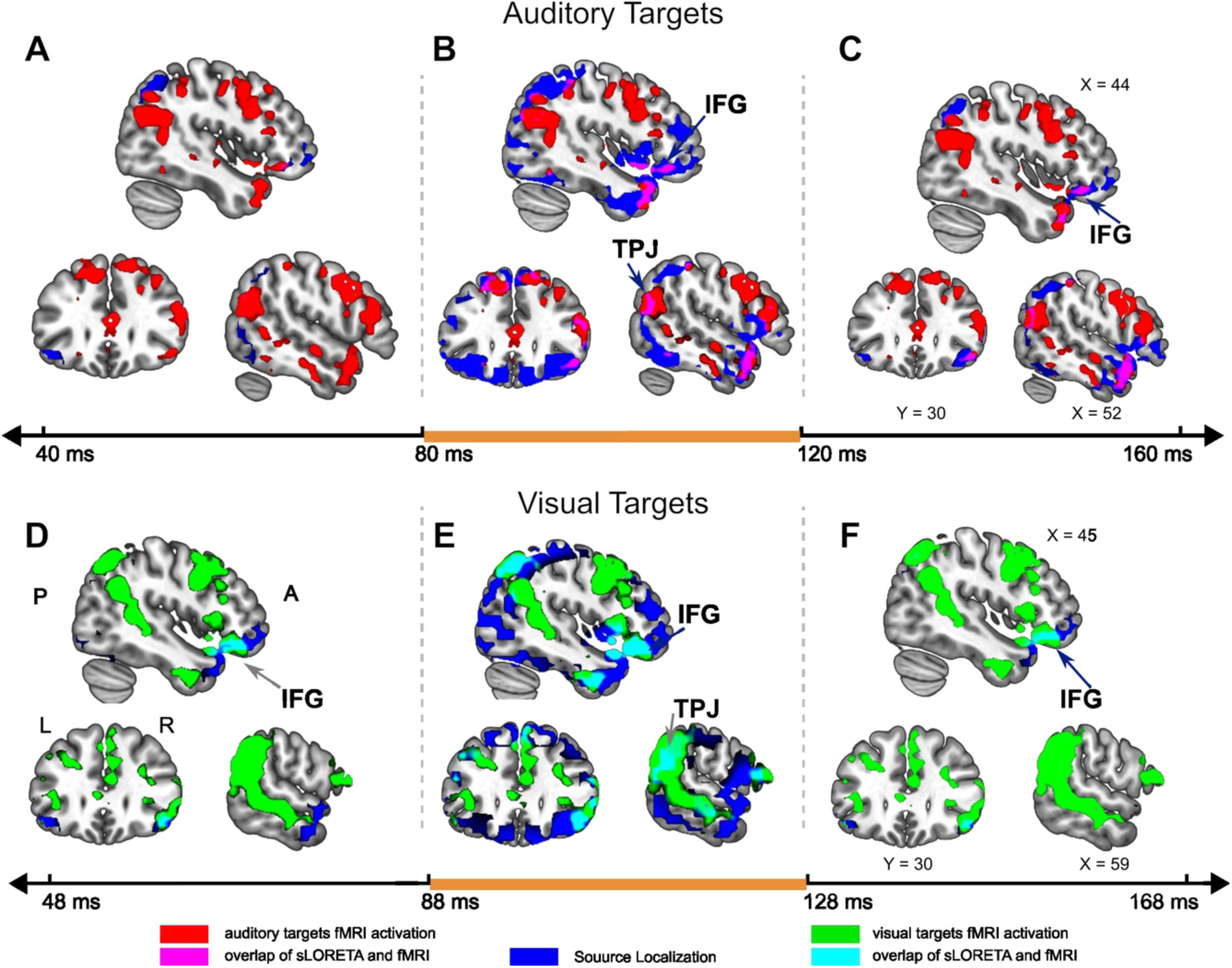
Time-resolved fMRI and source localization activation overlap. Top Row (Auditory Targets): **(A–C)** shows brain activation maps for auditory targets (unexpected > expected) at different time intervals following target onset. **(A)** At 40 ms post-stimulus, initial activations are observed primarily in the superior temporal and frontal regions. **(B**) At 80 ms, overlap spreads, with significant engagement in the inferior frontal gyrus (IFG) and temporoparietal junction (TPJ). **(C**) At 120 ms, the overlap is limited to right IFG. Red shading represents areas of fMRI activation, blue shading represents source localization using sLORETA, and purple areas indicate overlap between these maps. Bottom Row (Visual Targets): (D–F) show brain activations for visual targets (unexpected > expected) **(D)** At 48 ms, initial activations overlap in the right IFG. **(E)** By 88 ms, the right TPJ and IFG show increased overlaps. **(F)** Between 128-168 ms, the overlap is limited to right IFG only. Green shading represents areas of fMRI activation for visual targets, blue for source localization, and cyan areas indicate overlap between sLORETA and fMRI maps.

#### Connectivity between VAN and sensory areas

In bottom-up sensory processing, the right inferior frontal gyrus (IFG) is essential for inhibitory control and reallocation of attention, especially when encountering novel or unexpected stimuli. It has been widely recognized for its role in inhibiting prepotent responses when unexpected or novel stimuli are encountered (Aron et al., 2003; Hampshire et al., 2010). The right TPJ is involved in attentional reorientation, particularly in detecting and responding to unexpected, novel, or salient stimuli (Geng & Mangun, 2011; Geng & Vossel, 2013). Together, the IFG and TPJ coordinate within VAN to support the process of disengagement from ongoing, goal-directed tasks (driven by the dorsal attention network) and reorient toward unexpected but potentially important events. Indeed, in our study, we observed that the right IFG and TPJ showed heightened activity for unexpected auditory and visual stimuli, suggesting a role of VAN associated with prediction violations across modalities. Research suggests that the VAN supports directing attention to a task relevant stimulus and interacts with sensory areas (Maurizio Corbetta & Gordon L. Shulman, 2002; Egner et al., 2010; Hopfinger et al., 2000; Li et al., 2012; Sabine Kastner, 1999; Sreenivasan Meyyappan, 2021). Even though univariate analysis, including our study, shows evidence for shifts in frontoparietal activity in response to unexpected stimuli, in support of this theory, the evidence remains indirect. What is the relation of this activity within the VAN regions and regions outside VAN? The beta-series correlation method is a functional connectivity analysis technique that leverages trial-by-trial variability to assess covariance in activity across distant brain regions, offering a more direct approach to study network interactions (Rissman et al., 2004; Sreenivasan Meyyappan, 2021).

To understand how IFG interacts with TPJ in cross-modal predictive processing, we analyzed functional connectivity between these regions under different conditions. During unexpected auditory targets, functional connectivity between the right IFG and TPJ was significantly stronger (p = 0.009, d = 0.36) compared to expected targets (d = 0.25), as shown in Figure 7. A similar pattern emerged for visual targets, with higher functional connectivity (p = 0.002, d = 0.38) for unexpected versus expected targets (d = 0.03), as shown in Figure 8. No such connectivity differences were observed between the left IFG and TPJ, for auditory (p = 0.8) or visual targets (p = 0.20), indicating lateralized sensitivity in the right VAN to unexpected sensory events. Since we also observed greater activation to unexpected targets in sensory areas (Table 5, Table 6), which are implicated to be lower order areas in the hierarchical system where prediction errors are generated (Kok et al., 2014), we tested the functional connectivity between them and areas in the VAN. In case of auditory modality, we found greater functional connectivity between left STG and right TPJ for unexpected (d = 0.38) compared to expected (d = 0.2) auditory targets (p = 0.02). We did not observe the same effect in the right STG, or between right IFG and STG. Along similar lines, for visual modality, we found greater connectivity between left Fusiform Gyrus (FG) and right TPJ for unexpected (d = 0.28, p = 0.009) compared to expected targets (d = −0.13). We did not find any connectivity between right FG and regions in VAN. This indicates a likely involvement of the left FG (only) while processing visual gratings.

**Figure 7.**
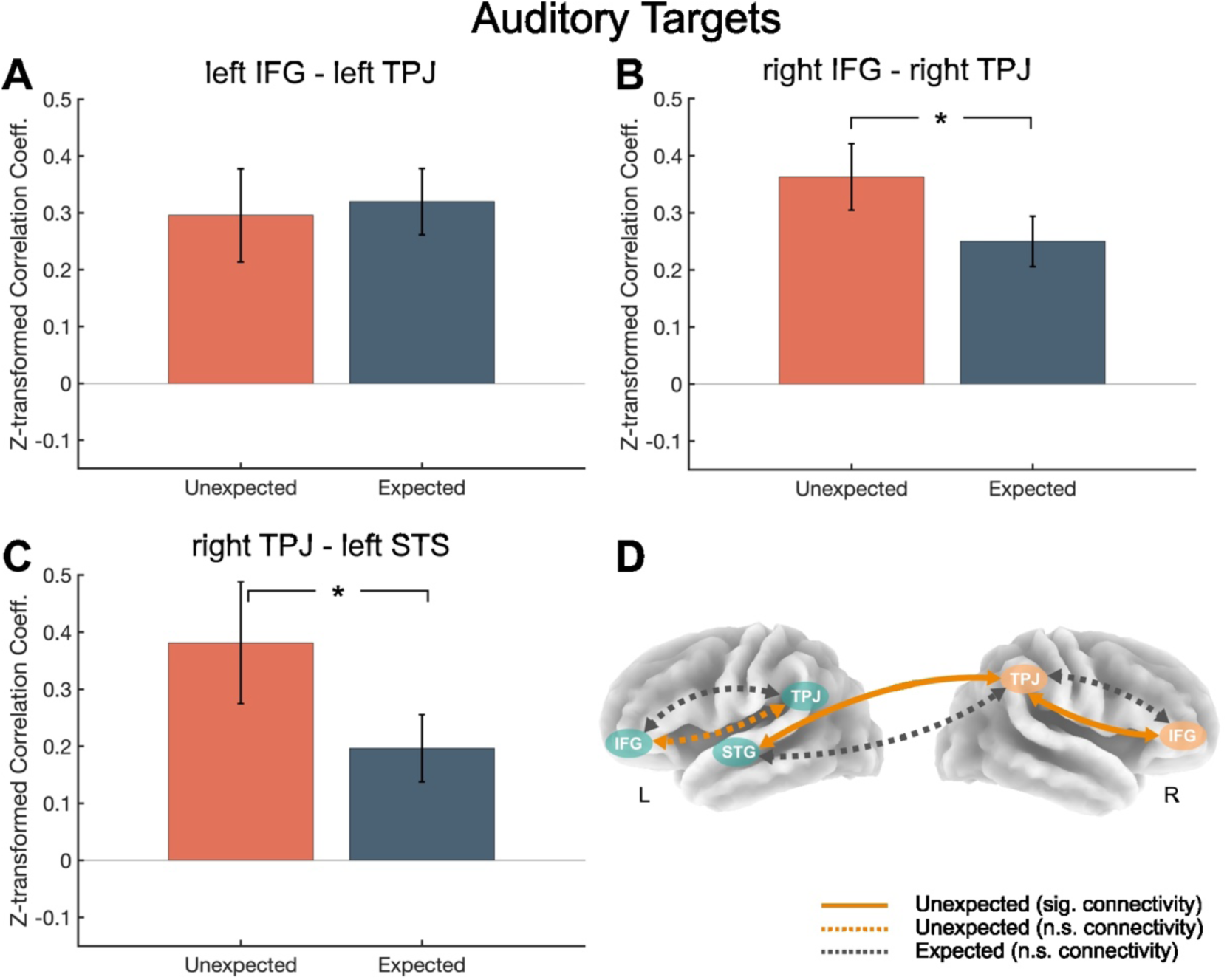
Functional connectivity analysis of auditory target processing between regions in VAN, comparing expected and unexpected auditory targets. **(A)** Connectivity between the left IFG and left TPJ shows no significant difference between unexpected and expected auditory targets. **(B)** Connectivity between the right IFG and right TPJ shows significantly higher connectivity for unexpected auditory targets compared to expected ones (*p = 0.009). **(C)** Connectivity between the right TPJ and left STG is also significantly higher for unexpected compared to expected auditory targets (*p = 0.02). **(D)** Functional connectivity is higher in the right-VAN during unexpected auditory targets. Further, right TPJ is functionally connected to the left STG. Solid orange lines indicate significant connectivity for unexpected auditory targets, while dashed lines represent non-significant connectivity. The color coding corresponds to expected and unexpected conditions across the regions: IFG = Inferior Frontal Gyrus, TPJ = Temporoparietal Junction, and STG = Superior Temporal Gyrus, VAN = Ventral Attention Network.

**Figure 8.**
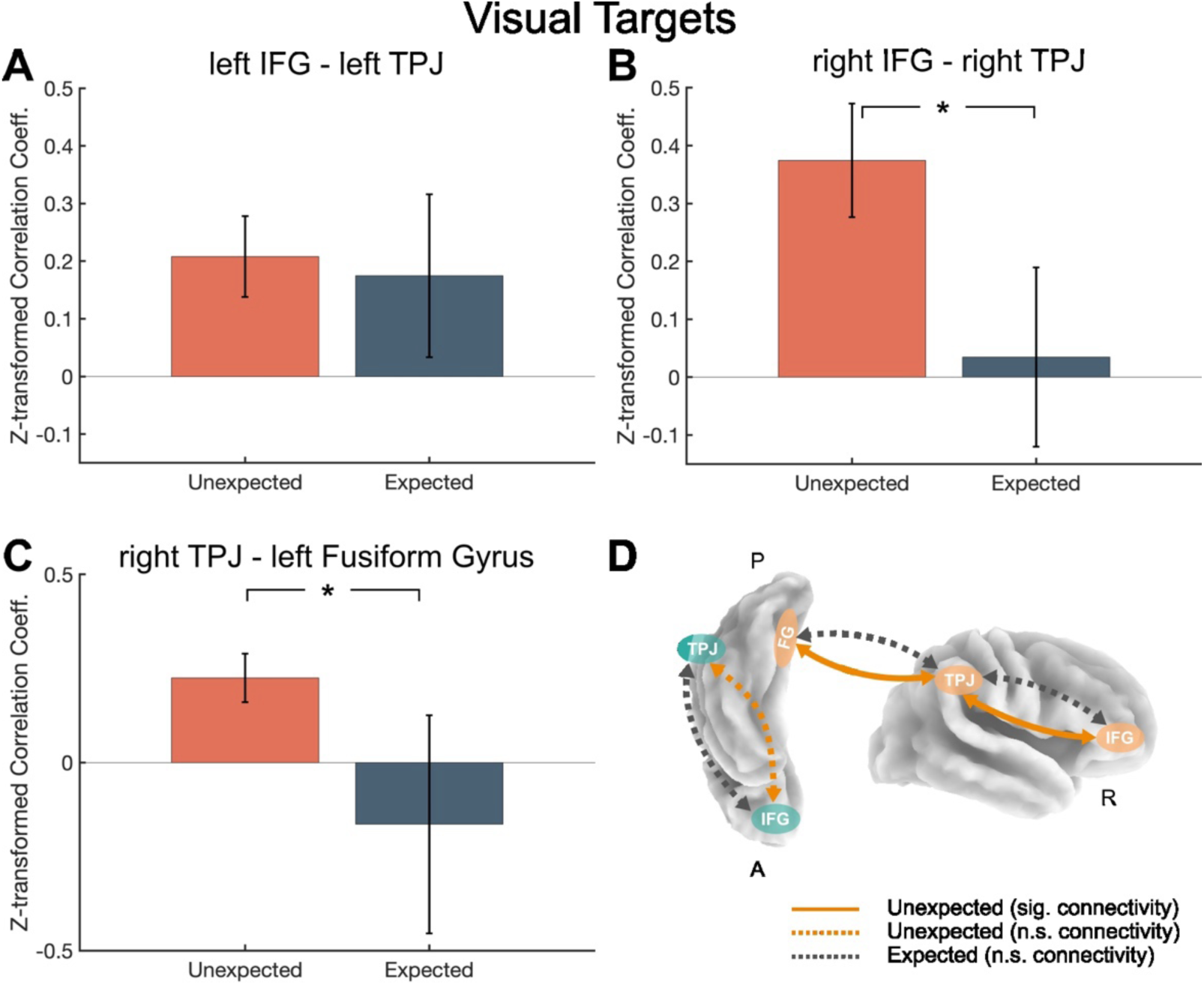
Functional connectivity analysis of visual stimuli processing within regions of the Ventral Attention Network (VAN), comparing expected and unexpected visual targets. **(A**) Connectivity between the left IFG and left TPJ shows no significant difference between expected and unexpected visual targets. **(B)** Connectivity between the right IFG and right TPJ is significantly higher for unexpected visual targets compared to expected visual targets (*p = 0.009). **(C)** Connectivity between the right TPJ and left FG is also significantly greater for unexpected visual targets compared to expected ones (*p = 0.01). **(D)** The schematic illustrates that functional connectivity in the right VAN is enhanced during unexpected visual targets, with additional connectivity between the right TPJ and left FG. Solid orange lines indicate significant connectivity for unexpected visual targets, while dashed lines represent non-significant connectivity. IFG = Inferior Frontal Gyrus, TPJ = Temporoparietal Junction, and FG = Fusiform Gyrus, VAN = Ventral Attention Network.

## Discussion

We investigated the neural effects of cross-modal expectations, focusing on the pivotal role of the Ventral Attention Network (VAN) in reorienting attention from one sensory modality to another when expectations are violated. Using an auditory-visual trial-by-trial paradigm with probabilistic cues, we induced expectations about the sensory modality of upcoming task-relevant stimuli. Across three experiments, several key findings emerged.

First, across both experiments (1 and 2), unexpected stimuli elicited more negative-going ERPs for both visual and auditory stimuli. Second, despite procedural variations, these early-latency ERP effects remained robust and consistent across experiments. Third, both sLORETA (ERP source localization) and fMRI confirmed right-lateralized activations in the temporoparietal junction (TPJ) and inferior frontal gyrus (IFG). Finally, increased connectivity between VAN regions (TPJ and IFG) and sensory processing areas was observed for unexpected stimuli. These findings are consistent with control processes originating in the VAN interacting with sensory structures to accomplish shifts of attention from attended to unattended sensory inputs. These control processes may involve attentional and predictive mechanisms, which would be expected to interact in circumstances like those in our cross-modal cuing paradigm. Further, this right-hemisphere bias in TPJ and IFG activation aligns with the VAN’s known lateralized functions in attentional reorientation (M. Corbetta & G. L. Shulman, 2002; Vossel et al., 2012) and perhaps predictive processing (Ficco et al., 2021).

### Early-Latency ERP Responses to Cross-Modal Expectation Violations

The findings from our study contribute significantly to our understanding of how sensory information across modalities is processed through the brain’s predictive framework, where predictions are continuously tested against incoming sensory inputs. This cross-modal expectation effect reflects the brain’s mechanism for efficiently managing sensory information by anticipating and validating stimuli across modalities. Mismatch studies, examining the mismatch negativity (MMN) component, demonstrate how the brain registers discrepancies between expected and actual input as prediction errors, traditionally in a single modality. Studies have shown differences in auditory stimuli evoked ERPs modulated by expectation peaking between 100-250 ms (Fitzgerald & Todd, 2020; Näätänen et al., 2007), over frontocentral scalp regions. Its visual counterpart, the vMMN emerges around 150 ms (Stefanics et al., 2015) at posterior scalp regions. Traditionally, this enhanced negativity is attributed to prediction errors that are generated in lower level sensory areas when conflict arises between priors and current sensory evidence (Garrido et al., 2009). Additionally, research on expectation-driven ERP suppression shows that predictable stimuli evoke attenuated neural responses, a phenomenon often referred to as “expectation suppression,” which has been widely observed in unisensory contexts but less often in cross-modal ones (Summerfield & Egner, 2016; Matthew F. Tang et al., 2018).

Across Experiments 1 and 2, we consistently observed a significant negative polarity ERP component within the preselected time window of 70–120 ms for auditory targets and 90– 150 ms for visual targets, with greater amplitude for unexpected compared to expected targets. Given our probabilistic cueing design, which is similar to the vast body of mismatch studies, this ERP component could be interpreted as an effect of surprise when stimulus occurs in a different modality (Egner et al., 2010; Richter & de Lange, 2019), or alternatively, an effect of expectation suppression (Feuerriegel et al., 2021). However, we argue that this is neither a pure surprise response nor simply a manifestation of expectation suppression. Instead, source localization and fMRI results provide converging evidence that this component may originate within the right-lateralized VAN, including the TPJ and IFG. This pattern is also observed during task-switching, where switching tasks involves similar brain areas and mechanisms (Czernochowski, 2011; Jamadar et al., 2015; Karayanidis & Jamadar, 2014; Li et al., 2012; Zhuo et al., 2021). For example, in a cued-task-switching study, Ling and colleagues (Li et al., 2012) reported a large switch-related central negative ERP component with sources in the dorsolateral prefrontal cortex including the MFG, IFG and posterior parietal cortex, suggesting an involvement of fronto-parietal networks in the task switching process. These areas are involved in mediating shifts of attention to goal-driven information and inhibition of response while detecting unexpected stimuli (Corbetta et al., 2008; Vossel et al., 2014). Interestingly,, using a bimodal stop-signal task in adults diagnosed with ADHD, Bekker and colleagues (Bekker et al., 2005) showed that successful stopping was associated with an enhanced early-latency negative ERP (around 100 ms) in medial frontocentral areas. Similarly, in our study, the brain must quickly disengage from the expected sensory modality and shift processing to the actual incoming stimulus, triggering an early-latency negative ERP component.

In our task, we encounter a unique case of mismatch processing where the brain must contend with more than just unmet predictions within a single sensory modality. Here, not only are predictions unmet in one modality—meaning the expected prediction errors are absent—but the incoming stimulus also arrives in a completely different modality (Foxe et al., 2005; Sidlauskaite et al., 2014). Some studies have pointed that omission responses can be associated with activity in units that deal with expectation rather than a pure reflection of error or surprisal (Wacongne et al., 2012; Walsh et al., 2020). Thus, rather than being solely a prediction error response or a manifestation of expectation suppression, the switch ERP observed in our study appears to reflect a broader mechanism that integrates perceptual expectation violations, task-switching, and attentional reorienting. The engagement of the right IFG and TPJ within the VAN suggests that the brain is not just detecting an unexpected stimulus but is actively resolving sensory conflict and reallocating attentional resources when the predicted modality is violated. While predictive coding frameworks (Arnal et al., 2011; Friston & Kiebel, 2009; M. F. Tang et al., 2018) offer a useful perspective suggesting that this ERP reflects an update in the brain’s predictive model—the observed effects align more closely with task-switching mechanisms.

### Neural circuitry of the Ventral Attention Network (VAN)

The engagement of right-lateralized IFG and TPJ shows an amodal role for VAN in processing cross-modal prediction violations, with each region playing a specialized function. It is interesting to note the temporal evolution of the VAN activation from the source localization - fMRI overlapping maps (Figure 6). Activity in the right IFG appears earlier, prior to the onset of the switch ERP effect. It has been speculated that the IFG sends a ‘stop signal’ through cortico-striatal-thalamic-cortical projections to inhibit a proponent response to upcoming stimuli (Bollinger et al., 2010; Chambers et al., 2009; Jahanshahi et al., 2015; Tomiyama et al., 2022). A meta-analysis of imaging studies found that individuals with OCD exhibit significantly reduced activation in striatal areas during inhibition tasks and, had reduced activation in prefrontal areas including the right IFG in switching tasks (Eng et al., 2015). Therefore, the early IFG activity in our study can be interpreted as an inhibitory mechanism when expectations are violated and existing priors need to be adapted. Following this, during the peak switch ERP window, both IFG and TPJ show significant activation. Activity in TPJ is more transient and tightly coupled with the peak switch ERP window in both auditory and visual conditions, suggesting a likely involvement of TPJ during the immediate reorientation phase. After this window, TPJ engagement declines, while IFG continues to show sustained activation. The temporal sequence in VAN, where IFG activation precedes TPJ, suggests a flow of information that supports adaptive response to cross-modal violations. IFG appears to initiate the process by inhibiting the irrelevant modality followed by interacting with TPJ for the reorientation. In fact, a task switching study combining fMRI and ERP measures found similar activations in the dorsolateral prefrontal areas during the preparation phase and another later ERP component in the posterior parietal areas after the onset of stimuli (Jamadar et al., 2010). This supports the idea that the TPJ is engaged at a later stage during evaluation of sensory events about prior expectations (Knight & Scabini, 1998; Vossel et al., 2014).

Finally, we examined the functional connectivity between regions in VAN in response to auditory and visual targets when they were expected compared to unexpected. Comparing auditory and visual target processing, both modalities demonstrated increased IFG-TPJ connectivity in response to unexpected stimuli (Figure 7 & Figure 8). Prior research has shown that expectations can modulate connectivity between prefrontal areas and TPJ by contextually updating top-down expectations or reorienting attention towards relevant tasks when external sensory evidence violates the current task set (Bressler et al., 2008; Chambon et al., 2017; Corbetta et al., 2008; Masina et al., 2022). We suggest that this functional link between IFG and TPJ during unexpected stimuli subserves for the switch and reorientation that is necessary to promote incoming sensory evidence which is relevant and inhibit any previously generated internal expectations. Beyond this shared pathway, we identified distinct connectivity patterns specific to each modality. For auditory targets, increased connectivity was observed with the left STG, whereas for visual targets, connectivity extended to the left FG. These modality-specific associations likely reflect the processing demands of sensory inputs that diverge from prior expectations (Johnston et al., 2017). The STG is known for its critical role in auditory processing (Yi et al., 2019) and has been implicated in encoding auditory prediction errors (Kim et al., 2024; Liu et al., 2023).

Similarly, a meta-analysis of prediction error studies reported increased activation in the left fusiform gyrus, which plays a key role in visual processing and the detection of prediction errors (D’Astolfo & Rief, 2017). This integration supports the notion of VAN working across sensory domains, continuously refining expectations based on incoming sensory inputs. The unique connectivity patterns observed between the TPJ and modality-specific regions reflect a coordinated feedback loop where VAN guides top-down modulation while enabling bottom-up adjustments based on sensory evidence. The greater connectivity between VAN and lower-order sensory areas for unexpected stimuli indicates that VAN’s role is not confined to attentional reorientation but extends to recalibrating sensory processing, enhancing perceptual sensitivity to unexpected but task-relevant stimuli (Rahnev et al., 2011).

### Rapid TPJ Activation Signals Cross-Modal Expectancy Violations

A well-established view holds that activity in TPJ is associated with the P300 ERP component, typically emerging between 250–400 ms post-stimulus (Bledowski et al., 2004; Menon et al., 1997; Soltani & Knight, 2000). This component is thought to reflect contextual updating, wherein the brain revises its internal model in response to unexpected but salient stimuli (Bledowski et al., 2004; Geng & Vossel, 2013). However, our cross-modal task-switching paradigm showed TPJ activity emerging much earlier, at around 100 ms—well before the conventional P300 window. This suggests that, beyond its role in later sensory evaluation, the TPJ is also recruited in an earlier phase of processing when attentional shifts between sensory modalities are required. The TPJ integrates inputs across multiple modalities and plays a key role in detecting and reorienting attention toward salient or unexpected stimuli (Geng & Vossel, 2013; Grundei et al., 2023; Shomstein et al., 2010). In the context of task switching, where cognitive control mechanisms dynamically adjust attentional priorities, the TPJ is well-positioned to detect deviations from expected sensory contingencies. When an expected relationship between modalities is violated, the TPJ is preferentially engaged, signaling the need for a rapid shift in processing priorities.

The observed TPJ activation during the early stages suggests that it may correspond to an early neural warning, rather than the downstream P300-related context update. In cognitive control frameworks, this early time window falls within the range of initial sensory and mismatch responses, which are crucial for rapid task adaptation. For instance, mismatch negativity (MMN), typically arises within 100–200 ms post-stimulus and is thought to reflect early prediction-error detection (Alain & Woods, 1997; Ritter et al., 1999). Similarly, studies on task-switching paradigms have demonstrated that early ERP components, including a frontocentral negativity occurring around 100 ms, signal the need for adjustments in response to changing task demands (Bekker et al., 2005; Mazaheri et al., 2010). Given the multisensory nature of our paradigm, the brain may have engaged the TPJ almost immediately—“flagging” the unexpected input and facilitating a shift in attentional resources toward the task-relevant sensory modality.

Neurophysiological evidence further supports this rapid TPJ engagement. Beauchamp and colleagues (Beauchamp et al., 2012) reported that unexpected visual stimulation elicited gamma-band TPJ activity within ∼100 ms, relating to its role in early-stage attentional reorienting. This indicate the TPJ can respond on a faster timescale when a salient change or mismatch is introduced, consistent with our findings. Rather than serving as an alternative to the P300, this early TPJ response may be an initial phase of a multi-step process, wherein a fast attentional reorientation primes subsequent modality adjustments (Naeije et al., 2016). In this framework, the TPJ may engage twice: an early transient burst signaling a shift in task-relevant processing and a later, sustained activation similar to a P300. For example, MEG studies of task-switching have shown that early neural markers of attentional reallocation predict the magnitude of later P300 responses in the TPJ (Naeije et al., 2016). Indeed, we observe a similar effect in our study (Figure S1), where the difference in magnitude of the switch ERP (unexpected – expected) correlates with the difference of the P300s. This suggests a strong relation between the initial prediction-error signal and later processes involved sensory evaluation and contextual updating.

In summary, the TPJ’s early activation in our study represents an automatic detection of a cross-modal expectancy violation—an attentional reorienting signal that precedes the P300. The observed early TPJ response likely serves as the first step in a hierarchical process, preparing the system for later-stage cognitive updating reflected in the P300. Rather than contradicting classical P300 interpretations, this finding complements them, providing more evidence in favor of the TPJ’s role in task-switching mechanisms that require rapid adaptation.

## Conclusion

Our findings emphasize the critical role of the Ventral Attention Network (VAN), in processing cross-modal predictions and expectation violations. We demonstrate how VAN supports our brain’s ability to reorient attention across sensory modalities. When unexpected stimulus with a different sensory modality appears violating our predictions (for example: auditory tones when expecting a visual grating), the VAN supports task-switching and suppressing prior expectations to effectively process the available stimuli. This coordinated response not only highlights the flexibility of the VAN in adaptive sensory processing but also broadens our understanding of predictive coding within a cross-modal context, where distinct sensory inputs demand a seamless integration of top-down expectations and bottom-up sensory feedback. These findings open pathways for future research on predictive processing across complex, real-world, multisensory environments, enhancing our understanding of how the brain continuously adapts to dynamic sensory inputs.

Looking forward, research should further investigate the temporal dynamics of VAN interactions with lower-order sensory cortices using methods with higher temporal precision, such as magnetoencephalography (MEG) (Solís-Vivanco et al., 2021), to map the exact timing of VAN engagement in cross-modal predictive processing. Additionally, it would be valuable to explore these interactions in more complex, naturalistic environments where cross-modal stimuli frequently conflict. Understanding how the VAN adapts in real-world contexts could enhance our knowledge of sensory integration and inform interventions for attentional disorders where predictive processing may be impaired.

## Acknowledgements

This work was supported by NIMH grant MH117991 to G.R.M and M.D, NSF grant BCS2318886 to G.R.M., and NSF grant BCS2318984 to G.R.M. We are grateful to Lee Miller, Joy Geng, Steve Luck, Maurizio Corbetta, Sabine Kastner, Sarah Shomstein and the members of our labs for their helpful comments and advice.

## Conflicts of interest

The authors declare no competing interests.

## Notes

### Competing Interest Statement

The authors have declared no competing interest.

## References

Alain, C., & Woods, D. L. (1997). Attention modulates auditory pattern memory as indexed by event-related brain potentials. Psychophysiology, 34(5), 534–546.

Altieri, N. (2014). Multisensory integration, learning, and the predictive coding hypothesis. Frontiers in Psychology, 5. 10.3389/fpsyg.2014.00257

Arnal, L. H., Wyart, V., & Giraud, A. L. (2011). Transitions in neural oscillations reflect prediction errors generated in audiovisual speech. Nat Neurosci, 14(6), 797–801. 10.1038/nn.2810

Aron, A. R., Fletcher, P. C., Bullmore, E. T., Sahakian, B. J., & Robbins, T. W. (2003). Stop-signal inhibition disrupted by damage to right inferior frontal gyrus in humans. Nat Neurosci, 6(2), 115–116. 10.1038/nn1003

Asplund, C. L., Todd, J. J., Snyder, A. P., & Marois, R. (2010). A central role for the lateral prefrontal cortex in goal-directed and stimulus-driven attention. Nature neuroscience, 13(4), 507–512. 10.1038/nn.2509

Auksztulewicz, R., & Friston, K. (2015). Attentional Enhancement of Auditory Mismatch Responses: a DCM/MEG Study. Cereb Cortex, 25(11), 4273–4283. 10.1093/cercor/bhu323

Bastos, Andre M., Usrey, W. M., Adams, Rick A., Mangun, George R., Fries, P., & Friston, Karl J. (2012). Canonical Microcircuits for Predictive Coding. Neuron, 76(4), 695–711. 10.1016/j.neuron.2012.10.038

Beauchamp, M. S., Sun, P., Baum, S. H., Tolias, A. S., & Yoshor, D. (2012). Electrocorticography links human temporoparietal junction to visual perception. Nature neuroscience, 15(7), 957–959.

Bekker, E. M., Overtoom, C. C. E., Kooij, J. J. S., Buitelaar, J. K., Verbaten, M. N., & Kenemans, J. L. (2005). Disentangling Deficits in Adults With Attention-Deficit/Hyperactivity Disorder. Archives of General Psychiatry, 62(10), 1129. 10.1001/archpsyc.62.10.1129

Bledowski, C., Prvulovic, D., Goebel, R., Zanella, F. E., & Linden, D. E. (2004). Attentional systems in target and distractor processing: a combined ERP and fMRI study. Neuroimage, 22(2), 530–540.

Bollinger, J., Rubens, M. T., Zanto, T. P., & Gazzaley, A. (2010). Expectation-Driven Changes in Cortical Functional Connectivity Influence Working Memory and Long-Term Memory Performance. The Journal of Neuroscience, 30(43), 14399–14410. 10.1523/jneurosci.1547-10.2010

Braga, R. M., Wilson, L. R., Sharp, D. J., Wise, R. J., & Leech, R. (2013). Separable networks for top-down attention to auditory non-spatial and visuospatial modalities. Neuroimage, 74, 77–86. 10.1016/j.neuroimage.2013.02.023

Bressler, S. L., Tang, W., Sylvester, C. M., Shulman, G. L., & Corbetta, M. (2008). Top-down control of human visual cortex by frontal and parietal cortex in anticipatory visual spatial attention. J Neurosci, 28(40), 10056–10061. 10.1523/JNEUROSCI.1776-08.2008

Čeponienė, R., Lepistö, T., Soininen, M., Aronen, E., Alku, P., & Näätänen, R. (2003). Event-related potentials associated with sound discrimination versus novelty detection in children. Psychophysiology, 41(1), 130–141. 10.1111/j.1469-8986.2003.00138.x

Chambers, C. D., Garavan, H., & Bellgrove, M. A. (2009). Insights into the neural basis of response inhibition from cognitive and clinical neuroscience. Neurosci Biobehav Rev, 33(5), 631–646. 10.1016/j.neubiorev.2008.08.016

Chambon, V., Domenech, P., Jacquet, P. O., Barbalat, G., Bouton, S., Pacherie, E., Koechlin, E., & Farrer, C. (2017). Neural coding of prior expectations in hierarchical intention inference. Scientific Reports, 7(1), 1278. 10.1038/s41598-017-01414-y

Corbetta, M., Patel, G., & Shulman, G. L. (2008). The reorienting system of the human brain: from environment to theory of mind. Neuron, 58(3), 306–324. 10.1016/j.neuron.2008.04.017

Corbetta, M., & Shulman, G. L. (2002). Control of goal-directed and stimulus-driven attention in the brain. In Nature Reviews Neuroscience (Vol. 3, pp. 201–215).

Corbetta, M., & Shulman, G. L. (2002). Control of goal-directed and stimulus-driven attention in the brain. Nat Rev Neurosci, 3(3), 201–215. 10.1038/nrn755

Czernochowski, D. (2011). ERP Evidence for Scarce Rule Representation in Older Adults Following Short, but Not Long Preparatory Intervals. Frontiers in Psychology, 2. 10.3389/fpsyg.2011.00221

Czigler, I., Balázs, L., & Pató, L. v. G. (2004). Visual change detection: event-related potentials are dependent on stimulus location in humans. Neuroscience Letters, 364(3), 149–153. 10.1016/j.neulet.2004.04.048

D’Astolfo, L., & Rief, W. (2017). Learning about Expectation Violation from Prediction Error Paradigms - A Meta-Analysis on Brain Processes Following a Prediction Error. Front Psychol, 8, 1253. 10.3389/fpsyg.2017.01253

Das, S., Yi, W., Ding, M., & Mangun, G. R. (2023). Optimizing cognitive neuroscience experiments for separating event-related fMRI BOLD responses in non-randomized alternating designs. Front Neuroimaging, 2, 1068616. 10.3389/fnimg.2023.1068616

Delorme, A., & Makeig, S. (2004). EEGLAB: an open source toolbox for analysis of single-trial EEG dynamics including independent component analysis. Journal of Neuroscience Methods, 134(1), 9–21. 10.1016/j.jneumeth.2003.10.009

den Ouden, H. E., Kok, P., & de Lange, F. P. (2012). How prediction errors shape perception, attention, and motivation. Front Psychol, 3, 548. 10.3389/fpsyg.2012.00548

Downar, J., Crawley, A. P., Mikulis, D. J., & Davis, K. D. (2000). A multimodal cortical network for the detection of changes in the sensory environment. Nature neuroscience, 3(3), 277–283. 10.1038/72991

Drisdelle, B. L., Aubin, S., & Jolicoeur, P. (2016). Dealing with ocular artifacts on lateralized ERPs in studies of visual-spatial attention and memory: ICA correction versus epoch rejection. Psychophysiology, 54(1), 83–99. 10.1111/psyp.12675

Egner, T., Monti, J. M., & Summerfield, C. (2010). Expectation and surprise determine neural population responses in the ventral visual stream. J Neurosci, 30(49), 16601–16608. 10.1523/JNEUROSCI.2770-10.2010

Eng, G. K., Sim, K., & Chen, S. H. (2015). Meta-analytic investigations of structural grey matter, executive domain-related functional activations, and white matter diffusivity in obsessive compulsive disorder: an integrative review. Neurosci Biobehav Rev, 52, 233–257. 10.1016/j.neubiorev.2015.03.002

Feuerriegel, D., Vogels, R., & Kovacs, G. (2021). Evaluating the evidence for expectation suppression in the visual system. Neurosci Biobehav Rev, 126, 368–381. 10.1016/j.neubiorev.2021.04.002

Ficco, L., Mancuso, L., Manuello, J., Teneggi, A., Liloia, D., Duca, S., Costa, T., Kovacs, G. Z., & Cauda, F. (2021). Disentangling predictive processing in the brain: a meta-analytic study in favour of a predictive network. Scientific Reports, 11(1). 10.1038/s41598-021-95603-5

Fitzgerald, K., & Todd, J. (2020). Making Sense of Mismatch Negativity. Frontiers in Psychiatry, 11. 10.3389/fpsyt.2020.00468

Foxe, J. J., Simpson, G. V., Ahlfors, S. P., & Saron, C. D. (2005). Biasing the brain’s attentional set: I. Cue driven deployments of intersensory selective attention. Experimental brain research, 166(3-4), 370–392. 10.1007/s00221-005-2378-7

Friston, K., & Kiebel, S. (2009). Predictive coding under the free-energy principle. Philosophical Transactions of the Royal Society B: Biological Sciences, 364(1521), 1211–1221. 10.1098/rstb.2008.0300

Friston, K. J., Holmes, A. P., Worsley, K. J., Poline, J.-P., Frith, C. D., & Frackowiak, R. S. J. (1994). Statistical parametric maps in functional imaging: A general linear approach. Human Brain Mapping, 2(4), 189–210. 10.1002/hbm.460020402

Gajewski, P. D., Ferdinand, N. K., Kray, J., & Falkenstein, M. (2018). Understanding sources of adult age differences in task switching: Evidence from behavioral and ERP studies. Neurosci Biobehav Rev, 92, 255–275. 10.1016/j.neubiorev.2018.05.029

Garrido, M. I., Kilner, J. M., Stephan, K. E., & Friston, K. J. (2009). The mismatch negativity: A review of underlying mechanisms. Clinical Neurophysiology, 120(3), 453–463. 10.1016/j.clinph.2008.11.029

Geng, J. J., & Mangun, G. R. (2011). Right temporoparietal junction activation by a salient contextual cue facilitates target discrimination. Neuroimage, 54(1), 594–601. 10.1016/j.neuroimage.2010.08.025

Geng, J. J., & Vossel, S. (2013). Re-evaluating the role of TPJ in attentional control: contextual updating? Neurosci Biobehav Rev, *37*(10 Pt 2), 2608-2620. 10.1016/j.neubiorev.2013.08.010

Grundei, M., Schmidt, T. T., & Blankenburg, F. (2023). A multimodal cortical network of sensory expectation violation revealed by fMRI. Human Brain Mapping, 44(17), 5871–5891.

Hall, M. G., Mattingley, J. B., & Dux, P. E. (2018). Electrophysiological correlates of incidentally learned expectations in human vision. Journal of neurophysiology, 119(4), 1461–1470. 10.1152/jn.00733.2017

Hampshire, A., Chamberlain, S. R., Monti, M. M., Duncan, J., & Owen, A. M. (2010). The role of the right inferior frontal gyrus: inhibition and attentional control. Neuroimage, 50(3), 1313–1319. 10.1016/j.neuroimage.2009.12.109

Hillyard, S. A., Vogel, E. K., & Luck, S. J. (1998). Sensory gain control (amplification) as a mechanism of selective attention: electrophysiological and neuroimaging evidence. Philosophical Transactions of the Royal Society of London. Series B: Biological Sciences, 353(1373), 1257–1270. 10.1098/rstb.1998.0281

Hopfinger, J. B., Buonocore, M. H., & Mangun, G. R. (2000). The neural mechanisms of top-down attentional control. In Nature neuroscience (Vol. 3, pp. 284–291).

Indovina, I., & Macaluso, E. (2007). Dissociation of stimulus relevance and saliency factors during shifts of visuospatial attention. Cereb Cortex, 17(7), 1701–1711. 10.1093/cercor/bhl081

Jahanshahi, M., Obeso, I., Rothwell, J. C., & Obeso, J. A. (2015). A fronto–striato– subthalamic–pallidal network for goal-directed and habitual inhibition. Nature Reviews Neuroscience, 16(12), 719–732. 10.1038/nrn4038

Jamadar, S., Hughes, M., Fulham, W. R., Michie, P. T., & Karayanidis, F. (2010). The spatial and temporal dynamics of anticipatory preparation and response inhibition in task-switching. Neuroimage, 51(1), 432–449. 10.1016/j.neuroimage.2010.01.090

Jamadar, S. D., Thienel, R., & Karayanidis, F. (2015). Task Switching Processes. In Brain Mapping (pp. 327–335). 10.1016/b978-0-12-397025-1.00250-5

Johnston, P., Robinson, J., Kokkinakis, A., Ridgeway, S., Simpson, M., Johnson, S., Kaufman, J., & Young, A. W. (2017). Temporal and spatial localization of prediction-error signals in the visual brain. Biological Psychology, 125, 45–57. 10.1016/j.biopsycho.2017.02.004

Karayanidis, F., & Jamadar, S. D. (2014). Event-related potentials reveal multiple components of proactive and reactive control in task switching. In Task switching and cognitive control. (pp. 200–236). Oxford University Press. 10.1093/acprof:osobl/9780199921959.003.0009

Kim, H. (2014). Involvement of the dorsal and ventral attention networks in oddball stimulus processing: A meta-analysis. Human Brain Mapping, 35(5), 2265–2284. 10.1002/hbm.22326

Kincade, J. M., Abrams, R. A., Astafiev, S. V., Shulman, G. L., & Corbetta, M. (2005). An Event-Related Functional Magnetic Resonance Imaging Study of Voluntary and Stimulus-Driven Orienting of Attention. The Journal of Neuroscience, 25(18), 4593–4604. 10.1523/jneurosci.0236-05.2005

Knight, R. T., & Scabini, D. (1998). Anatomic bases of event-related potentials and their relationship to novelty detection in humans. J Clin Neurophysiol, 15(1), 3–13. 10.1097/00004691-199801000-00003

Kok, P., Failing, M. F., & de Lange, F. P. (2014). Prior expectations evoke stimulus templates in the primary visual cortex. J Cogn Neurosci, 26(7), 1546–1554. 10.1162/jocn_a_00562

Kok, P., Rahnev, D., Jehee, J. F., Lau, H. C., & de Lange, F. P. (2012). Attention reverses the effect of prediction in silencing sensory signals. Cereb Cortex, 22(9), 2197–2206. 10.1093/cercor/bhr310

Li, L., Wang, M., Zhao, Q.-J., & Fogelson, N. (2012). Neural Mechanisms Underlying the Cost of Task Switching: An ERP Study. PLoS One, 7(7), e42233. 10.1371/journal.pone.0042233

Masina, F., Pezzetta, R., Lago, S., Mantini, D., Scarpazza, C., & Arcara, G. (2022). Disconnection from prediction: A systematic review on the role of right temporoparietal junction in aberrant predictive processing. Neuroscience & Biobehavioral Reviews, 138, 104713. 10.1016/j.neubiorev.2022.104713

Mayer, A., Schwiedrzik, C. M., Wibral, M., Singer, W., & Melloni, L. (2016). Expecting to See a Letter: Alpha Oscillations as Carriers of Top-Down Sensory Predictions. Cerebral Cortex, 26(7), 3146–3160. 10.1093/cercor/bhv146

Mazaheri, A., Coffey-Corina, S., Mangun, G. R., Bekker, E. M., Berry, A. S., & Corbett, B. A. (2010). Functional disconnection of frontal cortex and visual cortex in attention-deficit/hyperactivity disorder. Biol Psychiatry, 67(7), 617–623. 10.1016/j.biopsych.2009.11.022

Mazziotta, J., Toga, A., Evans, A., Fox, P., Lancaster, J., Zilles, K., Woods, R., Paus, T., Simpson, G., Pike, B., Holmes, C., Collins, L., Thompson, P., Macdonald, D., Iacoboni, M., Schormann, T., Amunts, K., Palomero-Gallagher, N., Geyer, S., … Mazoyer, B. (2001). A probabilistic atlas and reference system for the human brain: International Consortium for Brain Mapping (ICBM). Philosophical Transactions of the Royal Society of London. Series B: Biological Sciences, 356(1412), 1293–1322. 10.1098/rstb.2001.0915

Menon, V., Ford, J. M., Lim, K. O., Glover, G. H., & Pfefferbaum, A. (1997). Combined event-related fMRI and EEG evidence for temporal—parietal cortex activation during target detection. Neuroreport, 8(14), 3029–3037.

Muller-Gass, A., Stelmack, R. M., & Campbell, K. B. (2006). The effect of visual task difficulty and attentional direction on the detection of acoustic change as indexed by the Mismatch Negativity. Brain Research, 1078(1), 112–130. 10.1016/j.brainres.2005.12.125

Näätänen, R., Paavilainen, P., Rinne, T., & Alho, K. (2007). The mismatch negativity (MMN) in basic research of central auditory processing: A review. Clinical Neurophysiology, 118(12), 2544–2590. 10.1016/j.clinph.2007.04.026

Näätänen, R., & Picton, T. (2007). The N1 Wave of the Human Electric and Magnetic Response to Sound: A Review and an Analysis of the Component Structure. Psychophysiology, 24(4), 375–425. 10.1111/j.1469-8986.1987.tb00311.x

Naeije, G., Vaulet, T., Wens, V., Marty, B., Goldman, S., & De Tiège, X. (2016). Multilevel cortical processing of somatosensory novelty: a magnetoencephalography study. Frontiers in Human Neuroscience, 10, 259.

Oostenveld, R., Fries, P., Maris, E., & Schoffelen, J.-M. (2011). FieldTrip: Open Source Software for Advanced Analysis of MEG, EEG, and Invasive Electrophysiological Data. Computational Intelligence and Neuroscience, 2011, 1–9. 10.1155/2011/156869

Pascual-Marqui, R. D. (2002). Standardized low-resolution brain electromagnetic tomography (sLORETA): technical details. Methods Find Exp Clin Pharmacol, 24 *Suppl D*, 5–12.

Picton, T. W., Hillyard, S. A., Krausz, H. I., & Galambos, R. (1974). Human auditory evoked potentials. I. Evaluation of components. Electroencephalogr Clin Neurophysiol, 36(2), 179–190. 10.1016/0013-4694(74)90155-2

Posner, M. I., Snyder, C. R., & Davidson, B. J. (1980). Attention and the detection of signals. Journal of experimental psychology: General, 109(2), 160.

Rahnev, D., Lau, H., & de Lange, F. P. (2011). Prior expectation modulates the interaction between sensory and prefrontal regions in the human brain. J Neurosci, 31(29), 10741–10748. 10.1523/JNEUROSCI.1478-11.2011

Rao, R. P. N., & Ballard, D. H. (1999). Predictive coding in the visual cortex: a functional interpretation of some extra-classical receptive-field effects. Nature neuroscience, 2(1), 79–87. 10.1038/4580

Ren, J., Huang, F., Zhou, Y., Zhuang, L., Xu, J., Gao, C., Qin, S., & Luo, J. (2020). The function of the hippocampus and middle temporal gyrus in forming new associations and concepts during the processing of novelty and usefulness features in creative designs. Neuroimage, 214, 116751. 10.1016/j.neuroimage.2020.116751

Richter, D., & de Lange, F. P. (2019). Statistical learning attenuates visual activity only for attended stimuli. Elife, 8. 10.7554/eLife.47869

Rissman, J., Gazzaley, A., & D’Esposito, M. (2004). Measuring functional connectivity during distinct stages of a cognitive task. Neuroimage, 23(2), 752–763. 10.1016/j.neuroimage.2004.06.035

Ritter, W., Sussman, E., Deacon, D., Cowan, N., & Vaughan, H. G. (1999). Two cognitive systems simultaneously prepared for opposite events. Psychophysiology, 36(6), 835–838.

Rolls, E. T., Huang, C.-C., Lin, C.-P., Feng, J., & Joliot, M. (2020). Automated anatomical labelling atlas 3. Neuroimage, 206, 116189. 10.1016/j.neuroimage.2019.116189

Rungratsameetaweemana, N., Itthipuripat, S., Salazar, A., & Serences, J. T. (2018). Expectations Do Not Alter Early Sensory Processing during Perceptual Decision-Making. The Journal of Neuroscience, 38(24), 5632–5648. 10.1523/jneurosci.3638-17.2018

Sabine Kastner, M. A. P., Peter De Weerd, Robert Desimone, and Leslie G. Ungerleider. (1999). Increased Activity in Human Visual Cortex during Directed Attention in the Absence of Visual Stimulation. Neuron, 22.

Samaha, J., Boutonnet, B., Postle, B. R., & Lupyan, G. (2018). Effects of meaningfulness on perception: Alpha-band oscillations carry perceptual expectations and influence early visual responses. Scientific Reports, 8(1). 10.1038/s41598-018-25093-5

On the role of space and time in auditory processing, 5 340–348 (2001).

Shomstein, S., Lee, J., & Behrmann, M. (2010). Top-down and bottom-up attentional guidance: investigating the role of the dorsal and ventral parietal cortices. Exp Brain Res, 206(2), 197–208. 10.1007/s00221-010-2326-z

Shulman, G. L., McAvoy, M. P., Cowan, M. C., Astafiev, S. V., Tansy, A. P., d’Avossa, G., & Corbetta, M. (2003). Quantitative Analysis of Attention and Detection Signals During Visual Search. Journal of neurophysiology, 90(5), 3384–3397. 10.1152/jn.00343.2003

Sidlauskaite, J., Wiersema, J. R., Roeyers, H., Krebs, R. M., Vassena, E., Fias, W., Brass, M., Achten, E., & Sonuga-Barke, E. (2014). Anticipatory processes in brain state switching—Evidence from a novel cued-switching task implicating default mode and salience networks. Neuroimage, 98, 359–365.

Solís-Vivanco, R., Jensen, O., & Bonnefond, M. (2021). New insights on the ventral attention network: Active suppression and involuntary recruitment during a bimodal task. In Human Brain Mapping (Vol. 42, pp. 1699–1713).

Soltani, M., & Knight, R. T. (2000). Neural origins of the P300. Critical Reviews™ in Neurobiology, 14(3-4).

Spence, C., & Santangelo, V. (2009). Capturing spatial attention with multisensory cues: a review. Hear Res, 258(1-2), 134–142. 10.1016/j.heares.2009.04.015

Sreenivasan Meyyappan, A. R., George R. Mangun, Mingzhou Ding. (2021). Role of Inferior Frontal Junction (IFJ) in the Control of Feature versus Spatial Attention.

Stefanics, G. B., Astikainen, P., & Czigler, I. N. (2015). Visual mismatch negativity (vMMN): a prediction error signal in the visual modality. Frontiers in Human Neuroscience, 8. 10.3389/fnhum.2014.01074

Stekelenburg, J. J., & Vroomen, J. (2015). Predictive coding of visual-auditory and motor-auditory events: An electrophysiological study. Brain Res, 1626, 88–96. 10.1016/j.brainres.2015.01.036

Summerfield, C., & De Lange, F. P. (2014). Expectation in perceptual decision making: neural and computational mechanisms. Nature Reviews Neuroscience, 15(11), 745–756. 10.1038/nrn3838

Summerfield, C., & Egner, T. (2016). Feature-Based Attention and Feature-Based Expectation. Trends in cognitive sciences, 20(6), 401–404. 10.1016/j.tics.2016.03.008

Summerfield, C., Egner, T., Greene, M., Koechlin, E., Mangels, J., & Hirsch, J. (2006). Predictive codes for forthcoming perception in the frontal cortex. Science, 314(5803), 1311–1314. 10.1126/science.1132028

Sussman, E. S., Bregman, A. S., & Lee, W. W. (2014). Effects of task-switching on neural representations of ambiguous sound input. Neuropsychologia, 64, 218–229. 10.1016/j.neuropsychologia.2014.09.039

Tang, M. F., Smout, C. A., Arabzadeh, E., & Mattingley, J. B. (2018). Prediction error and repetition suppression have distinct effects on neural representations of visual information. Elife, 7, e33123. 10.7554/eLife.33123

Tang, M. F., Smout, C. A., Arabzadeh, E., & Mattingley, J. B. (2018). Prediction error and repetition suppression have distinct effects on neural representations of visual information. Elife, 7. 10.7554/eLife.33123

Thut, G., Nietzel, A., Brandt, S. A., & Pascual-Leone, A. (2006). Alpha-band electroencephalographic activity over occipital cortex indexes visuospatial attention bias and predicts visual target detection. J Neurosci, 26(37), 9494–9502. 10.1523/JNEUROSCI.0875-06.2006

Todorovic, A., & de Lange, F. P. (2012). Repetition suppression and expectation suppression are dissociable in time in early auditory evoked fields. J Neurosci, 32(39), 13389–13395. 10.1523/JNEUROSCI.2227-12.2012

Tomiyama, H., Murayama, K., Nemoto, K., Tomita, M., Hasuzawa, S., Mizobe, T., Kato, K., Ohno, A., Tsuruta, S., Togao, O., Hiwatashi, A., & Nakao, T. (2022). Increased functional connectivity between presupplementary motor area and inferior frontal gyrus associated with the ability of motor response inhibition in obsessive– compulsive disorder. Human Brain Mapping, 43(3), 974–984. 10.1002/hbm.25699

van Belle, J., Vink, M., Durston, S., & Zandbelt, B. B. (2014). Common and unique neural networks for proactive and reactive response inhibition revealed by independent component analysis of functional MRI data. Neuroimage, 103, 65–74. 10.1016/j.neuroimage.2014.09.014

Vogel, E. K., & Luck, S. J. (2000). The visual N1 component as an index of a discrimination process. Psychophysiology, 37(2), 190–203. https://www.ncbi.nlm.nih.gov/pubmed/10731769

Vossel, S., Geng, J. J., & Fink, G. R. (2014). Dorsal and Ventral Attention Systems. The Neuroscientist, 20(2), 150–159. 10.1177/1073858413494269

Vossel, S., Weidner, R., Driver, J., Friston, K. J., & Fink, G. R. (2012). Deconstructing the architecture of dorsal and ventral attention systems with dynamic causal modeling. J Neurosci, 32(31), 10637–10648. 10.1523/JNEUROSCI.0414-12.2012 32/31/10637 [pii]

Wacongne, C., Changeux, J. P., & Dehaene, S. (2012). A neuronal model of predictive coding accounting for the mismatch negativity. J Neurosci, 32(11), 3665–3678. 10.1523/jneurosci.5003-11.2012

Walsh, K. S., Mcgovern, D. P., Clark, A., & O’Connell, R. G. (2020). Evaluating the neurophysiological evidence for predictive processing as a model of perception. Annals of the New York Academy of Sciences, 1464(1), 242–268. 10.1111/nyas.14321

Weiss, Y., Cweigenberg, H. G., & Booth, J. R. (2018). Neural specialization of phonological and semantic processing in young children. Human Brain Mapping, 39(11), 4334–4348. 10.1002/hbm.24274

Yi, H. G., Leonard, M. K., & Chang, E. F. (2019). The Encoding of Speech Sounds in the Superior Temporal Gyrus. Neuron, 102(6), 1096–1110. 10.1016/j.neuron.2019.04.023

Zhuo, B., Zhu, M., Cao, B., & Li, F. (2021). More change in task repetition, less cost in task switching: Behavioral and event-related potential evidence. Eur J Neurosci, 53(8), 2553–2566. 10.1111/ejn.15113

