## Supplementary Figure 1 for "The Ventral Attention Network Mediates Attentional Reorienting to Cross-Modal Expectancy Violations: Evidence from EEG and fMRI"

### Supplementary Figures

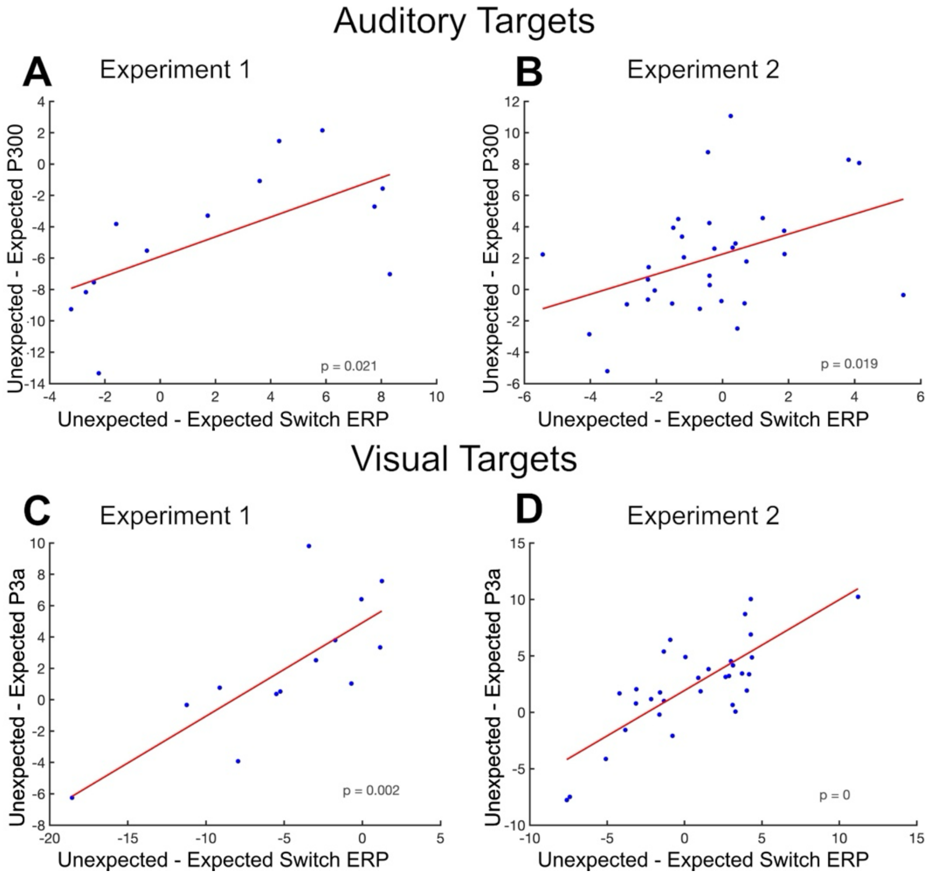


*Figure S 1* ***Correlation between P300/P3a and the switch ERP effect for unexpected – expected auditory (A-B) and visual (C-D) targets across Experiment 1 (A,C) and 2 (B,D).***
